# Enhanced translation expands the endo-lysosome size and promotes antigen presentation during phagocyte activation

**DOI:** 10.1101/260257

**Authors:** Victoria E. B. Hipolito, Jacqueline A. Diaz, Kristofferson V. Tandoc, Christian Oertlin, Johannes Ristau, Neha Chauhan, Amra Saric, Shannon Mclaughlan, Ola Larsson, Ivan Topisirovic, Roberto J. Botelho

## Abstract

The mechanisms that govern organelle adaptation and remodeling remain poorly defined. The endo-lysosomal system degrades cargo from various routes including endocytosis, phagocytosis and autophagy. For phagocytes, endosomes and lysosomes (endo-lysosomes) are kingpin organelles since they are essential to kill pathogens and process and present antigens. During phagocyte activation, endo-lysosomes undergo a morphological transformation, going from a collection of dozens of globular structures to a tubular network in a process that requires the phosphatidylinositol-3-kinase-AKT-mTOR signalling pathway. Here, we show that the endo-lysosomal system undergoes an expansion in volume and holding capacity during phagocyte activation within 2 h of LPS stimulation. Endo-lysosomal expansion was paralleled by an increase in lysosomal protein levels, but this was unexpectedly largely independent of TFEB and TFE3 transcription factors, known to scale up lysosome biogenesis. Instead, we demonstrate a hitherto unappreciated mechanism of acute organelle expansion via mTORC1-dependent increase in translation, which appears to be mediated by both, S6Ks and 4E-BPs. Moreover, we show that stimulation of RAW cells with LPS alters translation of a subset but not all of mRNAs encoding endo-lysosomal proteins, thereby suggesting that endo-lysosome expansion is accompanied by functional remodelling. Importantly, mTORC1-dependent increase in translation activity was necessary for efficient and rapid antigen presentation by dendritic cells. Collectively, we identified a previously unknown and functionally relevant mechanism for endo-lysosome expansion that relies on mTORC1-dependent translation to stimulate endo-lysosome biogenesis in response to an infection signal.

## Introduction

Eukaryotic cells compartmentalize a wide-range of biochemical functions within membrane-bound organelles such as the endoplasmic reticulum, peroxisomes, endosomes, and lysosomes. Of these, endosomes and lysosomes form the endo-lysosomal pathway, which receives, sorts, and traffics a multitude of endocytic and biosynthetic cargoes to either recycle or degrade. Typically, early and late endosomes are thought of as sorting stations, while lysosomes enable degradation and salvage of amino acids and other building units for cellular use. Yet, a more accurate view is that endosomes and lysosomes form a spectrum of heterogeneous tubulo-vesicular compartments, rather than defined populations [1, 2]. Indeed, late endosomes and lysosomes fuse to form hybrid endo-lysosomes, where degradation is thought to ensue [2]. Here, we refer to these structures as endo-lysosomes. Overall, eukaryotic organelles can exist in disparate morphologies ranging from individual vesicular organelles, stacks of flattened membrane sacs, to a continuous membrane reticulum, and can vary greatly in number, size, and activity. Importantly, cells can adapt organellar properties in response to a variety of intrinsic and extrinsic stimuli that alter the functional needs of cells [3–7]. Yet, how cells mold organellar properties in response to their differentiation state and/or change in their environment remains an outstanding question in cell biology.

Immune cells like macrophages and dendritic cells are plastic cells inasmuch as they can adopt “resting”, inflammatory, and anti-inflammatory states that differ in their gene expression profile, metabolism, secretory pathway activity, and endo-lysosomal membrane system [8–12]. With respect to the endo-lysosomal system, mature dendritic cells abate the degradative capacity of their endo-lysosomal system to help preserve antigenic peptides for presentation to adaptive immune cells [13]. On the other hand, macrophages enhance their lysosomal degradative power after phagocytosis to enhance bacterial killing [14]. Another example of endo-lysosomal remodelling occurs during lipopolysaccharide (LPS)-activation of macrophages and dendritic cells, which transform the endo-lysosomal system from a collection of dozens of individual globular organelles into a striking tubular network [12,15,16]. These tubules are positive for various endo-lysosomal markers such as LAMP1, CD63, Arl8b, Rab7, RILP, and in dendritic cells, they also comprise MHC-II, responsible for antigen presentation [15–17]. This reorganization requires downstream TLR4 signals including the phosphatidylinositol 3-kinase-AKT-mTOR axis, which may interface with Rab7 and Arl8b GTPases to control lysosome association with microtubule-motor proteins [16, 17]. These motors then help distort and tubulate endo-lysosomes on microtubule tracks [15,18,19]. While tubulation is associated with retention of pinocytic cargo, exchange of phagosomal cargo, and possibly antigen presentation [20–24], it is not presently known how endo-lysosome tubulation helps phagocytes perform their function in response to LPS and other stimulants.

Lysosomes also serve as signaling platforms to sense the metabolic and nutrient state of the cell [25–28]. For instance, a protein network involving the V-ATPase, ragulator and Rag GTPases sense high levels of amino acids within lysosomes to activate mTORC1 on the lysosome surface [29–34]. Active mTORC1 then phosphorylates various downstream targets to stimulate anabolic pathways, including mRNA translation [35–37]. In part, mTORC1 promotes translation by phosphorylating and activating the S6 kinases (S6Ks), which then act on multiple targets to modulate translation initiation, elongation and ribosome biogenesis [38–41]. This is coordinated by mTORC1-dependent phosphorylation of 4E-BPs which leads to their dissociation from eIF4E, thus allowing the eIF4F complex assembly and the recruitment of the ribosome to the mRNA [39,41,42]. Importantly, mTORC1-driven anabolic pathways are also coordinated with mTORC1-mediated repression of catabolic processes including autophagy. This is in large part achieved by phosphorylating ULK1, an initiator of autophagy, and inhibiting the transcription factor TFEB, which can up-regulate expression of lysosomal genes [43–46]. Inactivation of mTORC1 (e.g. during starvation) initiates autophagy, which is paralleled by a boost in lysosomal gene expression, and lysosomal activity to augment macromolecular turnover and help replenish the nutrients [45, 46]. mTORC1 also plays roles beyond coordinating anabolism and catabolism in the context of nutrient sensing. In macrophages and dendritic cells, mTORC1 activity is increased during Toll-like receptor ligand binding (e.g. LPS), *Salmonella* invasion, *Mycobacteria* infection, phagocytosis, and by inflammasome activators [14,17,47–51]. Though the functional consequences of increased mTORC1 activity in immune cells are not always clear, increased mTORC1 activity can lead to augmented protein synthesis and suppressed autophagy whereby both of these processes are thought to be required for stress resolution and cell survival [49,51,52].

Herein, we set out to further dissect the mechanisms underlying the reorganization of the endo-lysosomal system in activated phagocytes. We discovered that phagocytes expand their endo-lysosomal volume and retention capacity within two hours of LPS-mediated activation relative to their resting counterparts. We demonstrated that this expansion depends on augmented protein synthesis, but that this seems independent of transcriptional mechanisms such as activation of TFEB and TFE3. Instead, LPS-driven endo-lysosomal expansion depends on altered translation controlled by mTORC1 and its effectors, S6Ks and 4E-BPs. Interestingly, LPS-mediated enhanced translation was critical for rapid and efficient antigen presentation and T cell activation by dendritic cells. Ultimately, we present evidence that LPS engages mTORC1-dependent translation to increase endo-lysosome size and holding capacity.

## Results

### Activation of macrophage and dendritic cells expands the endo-lysosomal volume

Activation of macrophages and dendritic cells elicits a remarkable transformation of the endo-lysosome morphology, converting these organelles from dozens of individual puncta into a tubular network [17,20–22]. Upon careful visual inspection, we speculated that this tubular endo-lysosomal network occupied a larger volume than endo-lysosomes in resting cells (Fig 1a). To corroborate this observation, we quantified the total endo-lysosome volume in LPS-activated and resting cells by employing image volumetric analysis [53, 54]. We first pre-labelled endo-lysosomes with a fluorescent fluid-phase marker (see materials and methods) and then exposed cells to LPS or vehicle-alone for 2 h to induce endo-lysosome tubulation. Pre-labelling cells prior to stimulation ensured that endo-lysosomes were equally loaded with the dye in both resting and activated cells. We and others previously showed that endocytic tracers label tubules positive for various endo-lysosomal markers including LAMP1, CD63, Rab7, and MHC-II in macrophages and dendritic cells [15–17,21]. We then employed live-cell spinning disc confocal microscopy to acquire z-stacks and undertake volumetric analysis. Using this methodology, we observed a significant increase in volume occupied by the fluorescent probe in LPS-stimulated RAW macrophages, bone marrow-derived macrophages (BMDM) and bone marrow-derived dendritic cells (BMDCs) relative to their resting counterparts (Fig 1b). This suggests that activated phagocytes have an expanded total endo-lysosomal volume relative to resting cells.

**Fig 1:**
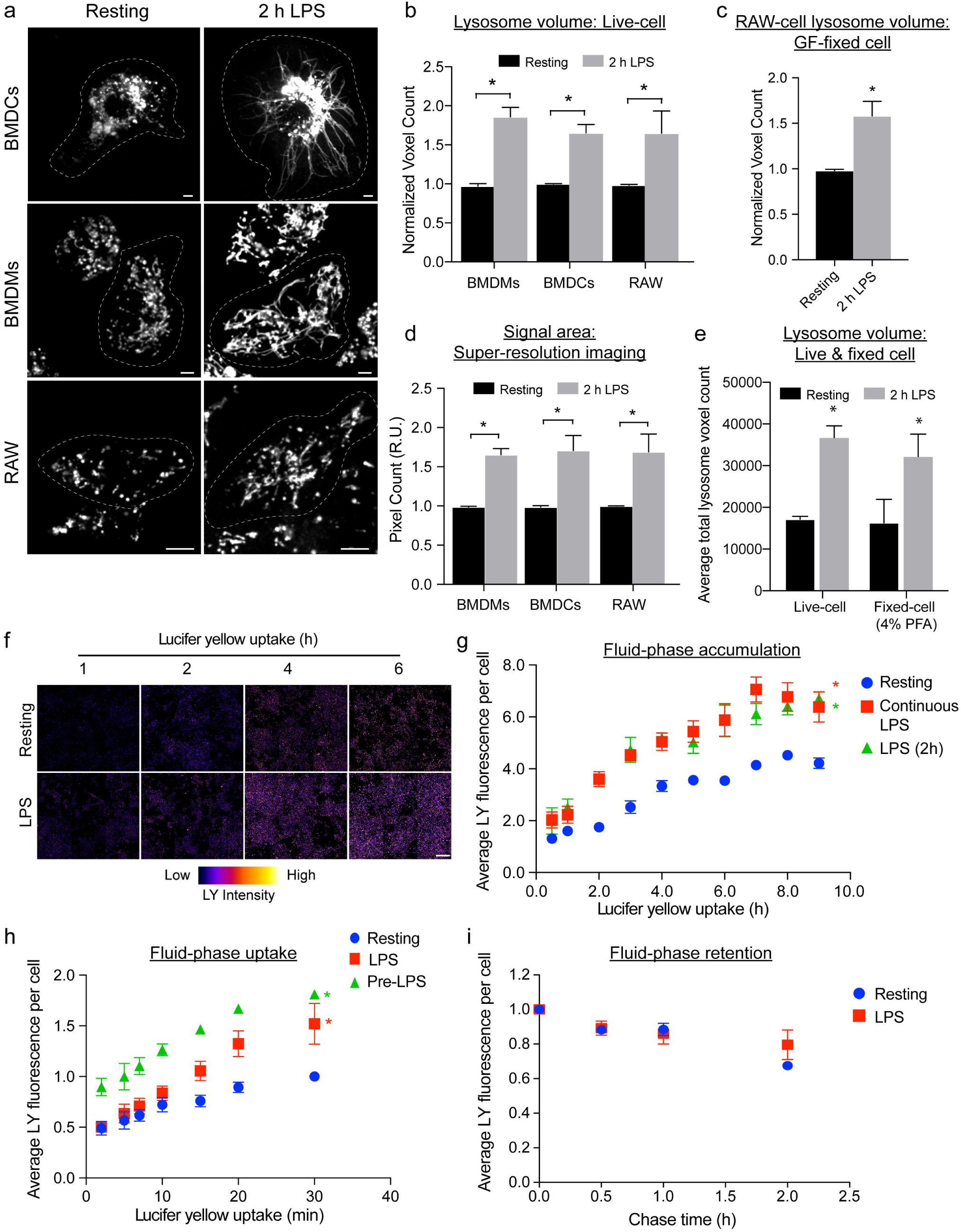
LPS-mediated activation of phagocytes augments lysosome volume and holding capacity. (a) Lysosomes in bone-marrow derived macrophages (BMDMs), bone-marrow derived dendritic cells (BMDCs), and in RAW macrophages before and after 2 h of LPS stimulation, the latter causing lysosome tubulation. Images were acquired by live-cell spinning disc confocal microscopy. Scale bar = 5 µm. (b) Relative lysosome volume between counterpart resting and LPS-treated phagocytes acquired by live-cell spinning disc confocal imaging. (c) Relative lysosome volume in resting and LPS-treated RAW macrophages fixed with a mixture of glutaraldehyde-formaldehyde (GF) to preserve tubules. (d) Relative lysosome area from the mid-section of resting and LPS-activated phagocytes using images acquired by SIM-enacted super-resolution microscopy. (e) Lysosome volume in resting and LPS-treated cells live or fixed with 4% PFA. (f) Image compilation of 6 representative fields in false-colour showing changes in intensity of Lucifer yellow (LY) acquired by endocytosis over the indicated time in resting primary macrophages or macrophages stimulated with LPS. Scale = 250 µm. Color scale: 0 – 4095 (low-high). (g) Accumulation of Lucifer yellow continuously endocytosed over indicated timeframe in resting, activated with LPS for 2 h or co-activated with LPS continuously. (h) Rate of pinocytosis of Lucifer yellow in primary macrophages treated as indicated. (i) Retention of Lucifer yellow in resting or LPS-treated primary macrophages after 0.5 h internalization and chase in probe-free medium over indicate times. All experiments were repeated at least three independent times. For b-e, data are based on 30-40 cells per condition per experiment and are shown as the mean ± SEM. Statistical analysis was performed using one-way ANOVA and unpaired post-hoc test, where the asterisk * indicates a significant increase in lysosome volume relative to resting phagocytes (p<0.05). For (g-i), fluorescence measurements were acquired by fluorimeter plate-imager. Data are shown as the mean ± SEM, where statistical analysis was performed using an Analysis of Covariance, whereby controlling for time as a continuous variable. An asterisk indicates a significant increase in Lucifer yellow for that series relative to resting phagocytes (p<0.05). See S1 Data for original data in Fig 1.

We previously demonstrated that in RAW macrophages, lysosome tubules were more motile than punctate lysosomes [15]. Thus, to exclude the possibility that the increase in endo-lysosome volume was due to a trailblazing effect during Z-stack image acquisition, we sought to estimate endo-lysosome volume in fixed cells. However, typical fixation protocols with 4% PFA causes tubular endo-lysosomes to collapse (S1a and S1b Figs). To circumvent this issue, we developed a fixation procedure that preserves endo-lysosome tubules in macrophages (S1a and S1b Figs). Re-applying volumetric analysis to fixed RAW cells, we still observed a significant increase in endo-lysosome volume in activated cells relative to resting phagocytes (Fig 1c). To then validate that the apparent increase in endo-lysosome volume triggered by LPS was not an artefact of imaging morphologically distinct objects we performed two tests. First, we employed structured illumination microscopy (SIM) in order to exclude the possibility that the limit of resolution of spinning disc confocal microscopy might cause an artefact when imaging morphologically-distinct endo-lysosomes [55]. Due to limitations of the available SIM system, we sampled three x-y planes centred at the mid-point of cells and quantified the area occupied by the fluid-phase marker (S1c Fig). This approach also revealed a significant increase in label area in activated RAW, primary macrophages, and BMDCs relative to their resting counterparts (Fig 1d). Second, we quantified the endo-lysosomal volume in LPS-treated cells fixed with 4% PFA. Herein, we took advantage that this treatment collapsed tubular endo-lysosomes into spheroid objects. As with other previous measurements, we observed higher endo-lysosome volume in fixed LPS-treated relative to resting macrophages (Fig 1e). Collectively, these data demonstrate that the endo-lysosome volume expands in response to macrophage and dendritic cell stimulation, concurrent with tubulation.

### Phagocyte activation increases endo-lysosomal holding capacity

An expanded endo-lysosomal volume may as a corollary lead to a boost in the storage capacity of endo-lysosomes. Hence, we assessed whether activated phagocytes have a higher endo-lysosomal holding capacity relative to resting cells by allowing cells to internalize fluorescent pinocytic tracers to saturation. Indeed, both primary and RAW macrophages pre-activated with LPS exhibited a significant increase in fluid-phase accumulation relative to their resting counterparts at each time point examined (Fig 1f and 1g; S2a Fig). We also observed that pre-activated primary macrophages displayed faster rates of pinocytic uptake relative to resting macrophages (Fig 1h). In fact, the rate of pinocytic uptake was augmented within 15 min of LPS exposure as indicated by macrophages concurrently undergoing pinocytosis and stimulation (Fig 1h). In comparison, we showed that resting and activated primary macrophages did not differ significantly in the rate of depletion of the pinocytic tracer (Fig 1i), suggesting that exocytosis rates were similar. RAW macrophages exhibited slightly different dynamics in that the rates of uptake and retention were similar between resting and LPS-stimulated cells (S2b and S2c Figs). Collectively, these data indicate that activated macrophages have a higher endo-lysosome holding capacity relative to resting macrophages.

Lastly, we questioned whether dendritic cells may benefit from an increase in endo-lysosomal volume since they are reported to arrest endocytosis after maturation [56, 57]. Of note, most reports examine dendritic cell function over 16 h post-stimulation, while recent work shows that mature cells can still endocytose extracellular cargo [58–60]. We show here that dendritic cells retained their pinocytic capacity up to 8 h post-activation, which fits the timeline of endo-lysosome reorganization and expansion observed here (S2d Fig). This suggests that expanding the endo-lysosomal volume within a few hours may help dendritic cells accumulate more pinocytic content including antigenic material. Overall, our observations are consistent with previous reports suggesting that tubulation in activated macrophages may aid in retaining fluid phase and that maturing dendritic cells continue to engulf extracellular material [20,58,59,61].

### Activated macrophages express higher levels of lysosomal proteins

Thus far, the data presented here suggest that phagocytes expand their endo-lysosomal volume and retention capacity within a couple of hours of activation. Though other mechanisms such as increased endosomal membrane influx may contribute to this, we postulated that endo-lysosomal biosynthesis may be a significant driver of endo-lysosome expansion during phagocyte activation. To address this hypothesis, we tracked the levels of seven major endo-lysosomal proteins as indicators of lysosome biogenesis; namely, we measured the levels of LAMP1, LAMP2, TRPML1, CD63, the V-ATPase subunits H and D, and cathepsin D by Western blotting in resting and activated primary macrophages. Specifically, we compared resting macrophages to those continuously exposed to LPS for 2 h or 6 h or for 2 h with LPS followed by a 4 h chase with no LPS. With the exception of cathepsin D, LPS induced the levels of all proteins approximately 2-fold when compared to resting macrophages (Fig 2a and 2b; S3 Fig). In addition, immunofluorescence staining for LAMP1 corroborates the increase in LAMP1 levels which was detected by Western blotting (Fig 2c and 2d). The increase in the levels of these lysosomal proteins was blunted by the translation elongation inhibitor, cycloheximide (Fig 2a and 2b). Finally, there was little change in the levels of proteins examined between resting cells or those treated with cycloheximide with or without LPS, suggesting similar turnover rates of these proteins (Fig 2a and 2b). Collectively, these data indicate that *de novo* protein synthesis, rather than lower protein turnover, augments the levels of lysosomal proteins in LPS-treated phagocytes (Fig 2a and 2b). Interestingly, cycloheximide blunted endo-lysosome tubulation and expansion in macrophages in response to LPS (Fig 2e and 2f), suggesting that *de novo* protein synthesis is required to remodel the endo-lysosome network during phagocyte activation. Overall, our data intimate that phagocytes boost protein synthesis to expand their endo-lysosomal system within 2 h of stimulation.

**Fig 2:**
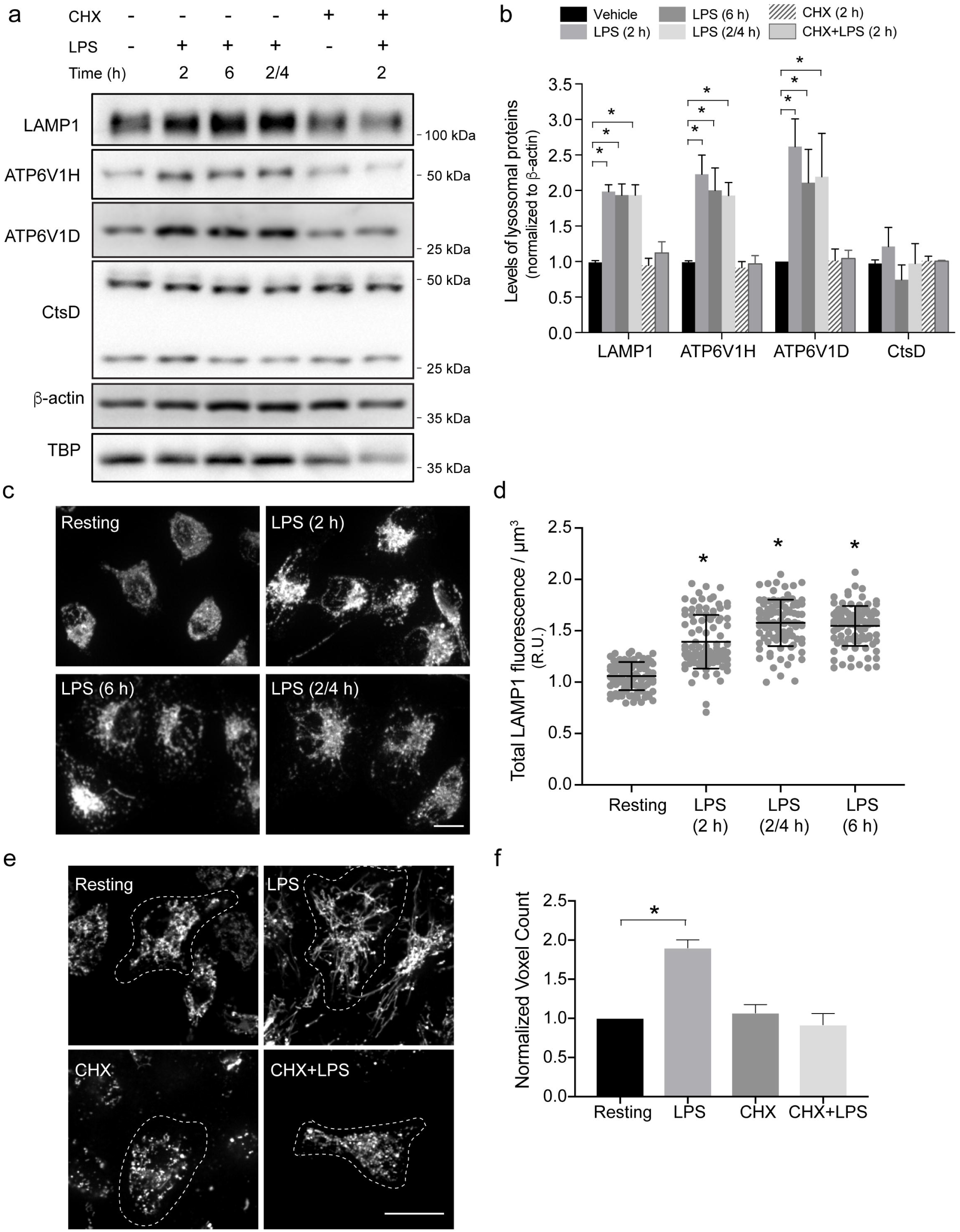
Lysosome remodelling requires protein biosynthesis. (a) Western blot analysis of whole cell lysates from resting primary macrophages or macrophages exposed to the indicated combinations and time of LPS and cycloheximide (CHX). (b) Quantification of Western blots showing the levels of LAMP1, cathepsin D (CtsD) and the V-ATPase V_1_ subunit H and D normalized to β-actin. Data are shown as the mean ± standard error of the mean from at least 3 independent experiments. For A and B, “2/4” indicates cells stimulated with 2 h of LPS, followed by a 4 h chase, whereas 2 and 6 h represent cells continuously exposed to LPS for those time periods. (c) Endogenous LAMP1-positive structures in resting and activated primary macrophages. (d) Quantification of total LAMP1 fluorescence levels in macrophages per μm^3^. (e) Live-cell spinning disc confocal micrographs of pre-labelled lysosomes in resting primary macrophages or those stimulated with LPS and/or cycloheximide. (f) Relative lysosome volume between resting primary macrophages and those exposed to specified conditions. Shown is the mean ± standard error of the mean from 30-40 cells for each condition and experiment, across at least 3 independent experiments. Scale bars = 5 µm Statistical analysis was done with ANOVA and unpaired post-hoc test. The asterisk * indicates a significant difference (p<0.05). For each figure with Western blots, see S1 Raw Images for original, unedited Western blots. See S2 Data for original data in Fig 2.

### Acute endo-lysosome expansion is not dependent on TFEB and TFE3

Our results suggest that biosynthesis plays a major role in LPS-induced endo-lysosome expansion in macrophages. Activation of TFEB and TFE3 transcription factors drives transcription of lysosomal genes thereby stimulating lysosome function under various stresses including starvation, phagocytosis, protein aggregation and macrophage activation [14,45,62–68]. Thus, we next investigated whether the observed endo-lysosome expansion was driven by TFEB- and TFE3-mediated transcriptional upregulation of lysosome genes.

To assess activation of TFEB and TFE3, we quantified their nuclear translocation by determining the nucleo-cytoplasmic ratio of endogenously expressed proteins by immunofluorescence [14, 69]. As expected [45, 46], resting cells exhibited mostly cytoplasmic TFEB and TFE3, whereas inhibition of mTOR for 1 h with torin1 caused both proteins to translocate into the nucleus (Fig 3a and 3b). Strikingly, while 2 h incubation with LPS was sufficient to induce endo-lysosome expansion, this did not trigger nuclear translocation of TFEB or TFE3 (Fig 3a and 3b). LPS triggered nuclear entry of these proteins only after 6 h of exposure (Fig 3a and 3b). These results are consistent with observations by Pastore *et al.*, who also observed delayed nuclear entry of these proteins in response to LPS-induced macrophage activation, which suggests that LPS stimulation of TFEB/TFE3 is indirect [66].

**Fig 3:**
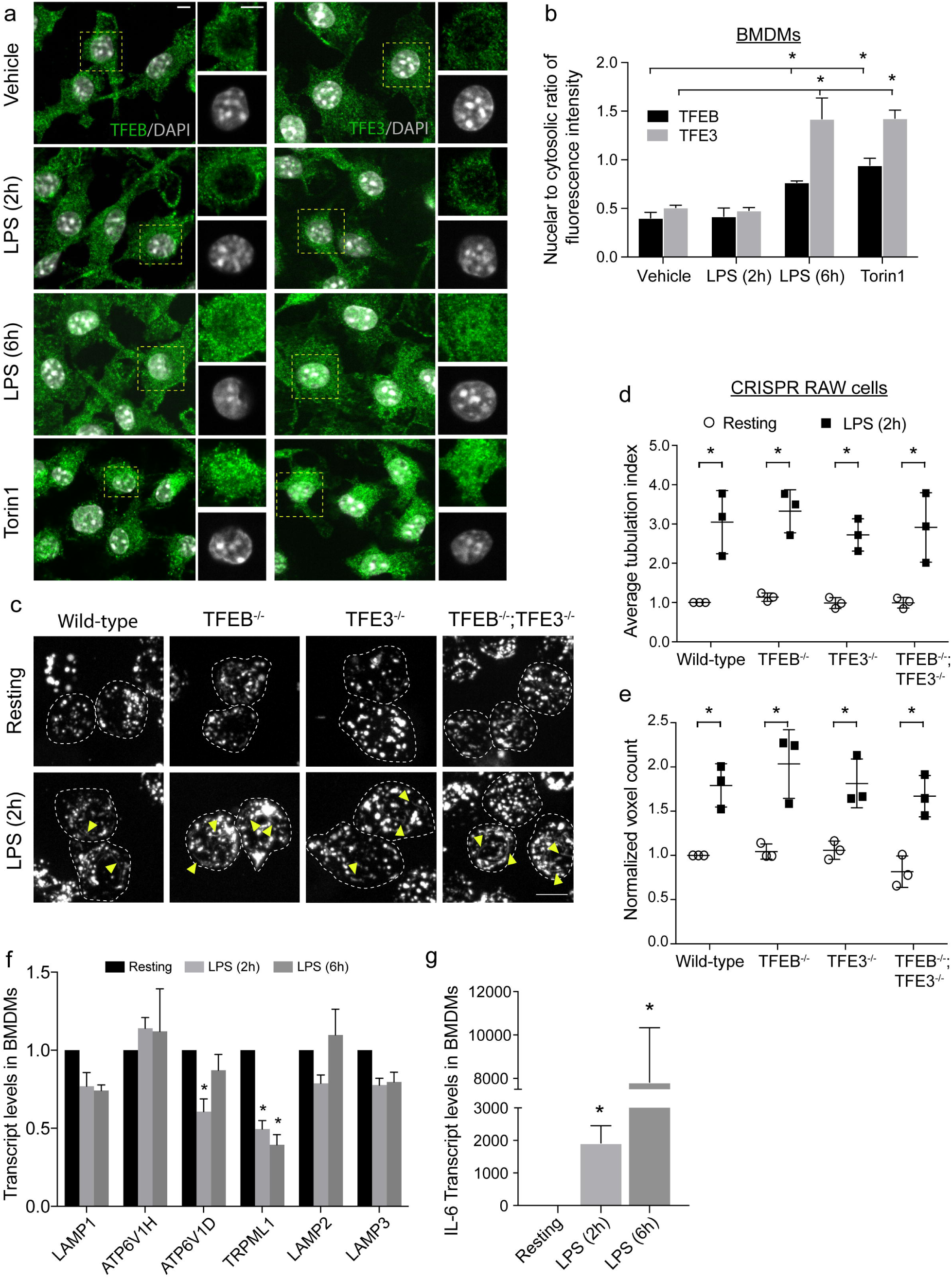
Lysosome remodelling is independent of TFEB and TFE3 activation. (a) TFEB and TFE3 subcellular localization in resting primary macrophages (vehicle) or those treated with LPS for 2 or 6 h, or with torin1. Green = TFEB or TFE3 immunofluorescence signal; white = nuclei stained with DAPI. Areas within dashed boxes are magnified as insets. (b) Nuclear to cytosolic ratio of TFEB or TFE3 fluorescence intensity. Shown is the mean ± standard error of the mean from 30-40 cells per condition per experiment across at least 3 independent experiments. (c) Lysosomes in wild-type, *tfeb^−/−^, tfe3*^−/−^ and *tfeb^−/−^ tfe3^−/−^* RAW strains before and after 2 h of LPS stimulation. Images were acquired by live-cell spinning disc confocal microscopy. Yellow arrowheads illustrate tubular lysosomes. (d) Average lysosome tubulation index in resting and LPS-activated strains. Shown is the mean ± standard error of the mean from 40-50 cells per condition, across three independent experiments. Lysosome tubules longer than 4 microns were scored, where tubulation index was determined following normalization to the average number of tubular lysosomes in resting wild-type cells for each experiment. (e) Relative lysosome volume between LPS-treated and resting counterpart RAW strains acquired by live-cell spinning disc confocal imaging. The average lysosomal voxel counts for LPS-activated strains were normalized to resting wild-type cells. Shown is the mean ± standard error of the mean from 30-40 cells per condition per experiment across three independent experiments. (f, g) Relative mRNA levels of select lysosomal genes (f) or interleukin-6 (g) in activated primary macrophages relative to Abt1 housekeeping gene and normalized against resting cells. Quantification was done with qRT-PCR by measuring the ÄÄCt as described in methods. Shown is the mean ± standard error of the mean from four independent experiments. All statistical analysis was done with ANOVA and unpaired post-hoc test. The asterisk * indicates a significant difference relative to resting condition (p<0.05). For (a) and (c), scale bar = 5 µm. See S3 Data for original data in Fig 3.

To further test whether TFEB does not play a role in endo-lysosome expansion during macrophage activation, we measured tubulation and endo-lysosome volume in RAW macrophages deleted for the genes encoding TFEB and/or TFE3 using CRISPR-based technology ([66], S4a and S4b Fig). Deletion of TFEB and TFE3 did not significantly affect LAMP1 protein levels under resting conditions (S4c and S4d Fig), nor trafficking of the fluid-phase marker we employed, as quantified by Mander’s coefficient for dextran-containing LAMP1 signal (S4e and S4f Fig). Moreover, both resting wild-type and deletion strains of RAW macrophages accumulated similar levels of the dextran probe after 1 h of uptake and 1 h chase (S4g Fig). Finally, TFEB and TFE3 status in the cell did not exert major influence on retention of the fluid-phase probe after 2 h of LPS exposure (S4h Fig). Collectively, these data suggest that TFEB and/or TFE3 have minimal impact on basal pinocytosis and basal biogenesis and trafficking to endo-lysosomes.

We next examined remodelling of endo-lysosomes by treating wild-type and TFEB and/or TFE3-deleted RAW cells with LPS for up to 2 h. Importantly, all three mutant cell lines exhibited an increase in endo-lysosome tubulation after 2 h of LPS treatment relative to the resting condition. This increase in endo-lysosome tubulation in cells depleted of TFEB and/or TFE3 was indistinguishable from that observed in wild-type RAW cells (Fig 3c and 3d). However, we do note that endo-lysosome tubulation in RAW cells is less pronounced than in primary macrophages and dendritic cells. Importantly, LPS-induced expansion of the total endo-lysosome volume was comparable between control and TFEB and/or TFE3-deleted cells (Fig 3e). These results suggest that TFEB and/or TFE3-dependent transcription-based programs are not required for lysosome expansion during acute macrophage activation.

Finally, to assess if other transcriptional mechanisms might be involved in LPS-induced lysosome expansion during the first couple of hours of activation, we measured mRNA levels encoding the six major endo-lysosomal proteins that increased in their levels in primary macrophages. We found that mRNA levels for LAMP1, LAMP2, CD63, TRPML1 and two V-ATPase subunits did not increase even after 6 h of LPS exposure (Fig 3f). In comparison, we observed a massive upregulation of interleukin-6 mRNA after LPS exposure (Fig 3g). Collectively, these data intimate that increased transcription does not explain increased levels of the corresponding proteins we previously observed in Fig 2 and S3 Fig. While it remains possible that transcriptional regulation of other mRNAs encoding endo-lysosome-related proteins contributes to endo-lysosome expansion, in particular during more prolonged stimulation, we speculated that post-transcriptional processes play a more pressing role in the LPS-mediated growth of the endo-lysosomal system.

### Endo-lysosome expansion depends on AKT and mTOR activity

Given that the levels of six major endo-lysosomal proteins, but not corresponding mRNAs were induced by LPS treatment, we next studied the role of translation in endo-lysosome expansion. Activated macrophages exhibit extensive metabolic reorganization, enhanced protein synthesis, selective translation of mRNAs encoding inflammatory proteins, and activation of unfolded protein response [8,10,11,49,70,71]. Consistently, LPS activates mTORC1 in macrophages, which not only stimulates mRNA translation, but is also necessary for endo-lysosome tubulation [17,36,37]. Thus, we tested whether mTOR activity is also necessary for enhanced endo-lysosome volume and holding capacity. Indeed, as suggested by others [72–74], both primary and RAW macrophages exhibited increased phosphorylation of mTORC1 substrates S6K and 4E-BP1 after exposure to LPS, which is blunted by torin1, an active-site mTOR inhibitor (S5a and S5c Fig). Moreover, consistent with our previous observations [17], endo-lysosome tubulation was suppressed upon inhibition of mTOR or AKT by torin1 and Akti, respectively (Fig 4a). Importantly, we now show that suppression of AKT and/or mTOR activity abrogates the LPS-induced expansion of the endo-lysosome volume (Fig 4a and 4b). Moreover, Akti and torin1 both prevented the increased in the levels of LAMP1, LAMP2, CD63, TRPML1 and V-ATPases elicited by LPS (Fig 4c and 4d; S3a and S3b Fig), suggesting that Akt-mTORC1 pathway is required for endo-lysosome expansion. In addition, mTOR inhibition also blunted the increase in the pinocytic holding capacity enticed by LPS treatment (Fig 4e). These effects are not likely due to autophagy induced by torin1 since Akti did not induce autophagy, but yet blocked lysosome expansion (S5d and S5e Fig). Collectively, these findings demonstrate that LPS-mediated stimulation of the Akt-mTOR pathway promotes expansion of the endo-lysosome system and retention capacity.

**Fig 4:**
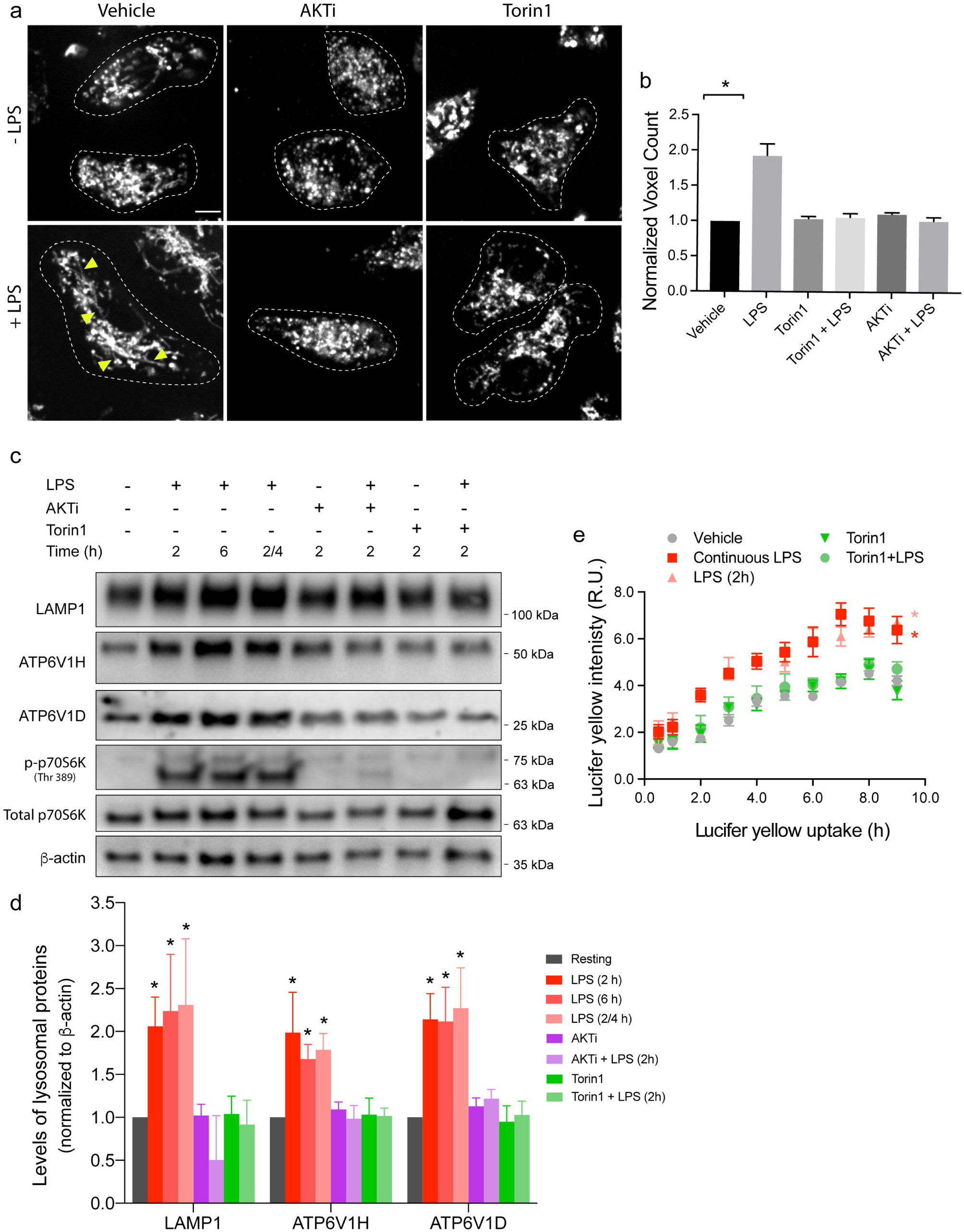
mTOR stimulates lysosome volume and holding capacity. (a) Lysosomes in primary macrophages were pre-treated with a vehicle (DMSO), Akti or torin1, followed by 2 h LPS stimulation where indicated. Images were acquired by live-cell spinning disc confocal microscopy. Scale bar = 5 µm. (b) Lysosome volume in primary macrophages treated as indicated and normalized to resting macrophages. Shown is the mean ± standard error of the mean from 30-40 cells per condition per experiment across three independent experiments. (c) Western blot analysis of whole cell lysates from resting primary macrophages or macrophages exposed to the indicated combinations and time of LPS, Torin1 and AKTi. (d) Quantification of Western blots showing the levels of LAMP1, cathepsin D (CtsD) and the V-ATPase V_1_ subunit H and D normalized to β-actin. Data are shown as the mean ± standard error of the mean from at least 3 independent experiments. For a and b, “2/4” indicates cells stimulated with 2 h of LPS, followed by a 4 h chase, whereas 2 and 6 h represent cells continuously exposed to LPS for those time periods. (e) Quantification of pinocytic capacity in macrophages treated as indicated. Shown is the mean ± standard error of the mean from four independent experiments. For b and d, data was statistically analysed with ANOVA and unpaired post-hoc test (*p<0.05). For e, data was statistically assessed using an Analysis of Covariance, whereby controlling for time as a continuous variable. An asterisk indicates a significant increase in Lucifer yellow for that series relative to resting phagocytes (*p<0.05). For each figure with Western blots, see S1 Raw Images for original, unedited Western blots. See S4 Data for original data in Fig 4.

### LPS triggers increased endo-lysosome synthesis via mTORC1 pathway

Given that mTOR is hyperactivated in LPS-exposed phagocytes and its activity is necessary for endo-lysosome expansion, we next tested whether LPS stimulates global protein synthesis in primary macrophages by employing the puromycylation assay. In this assay, cells are pulsed with puromycin, which is covalently added to growing peptides by the ribosome. Puromycin-tagged peptides can then be quantified with anti-puromycin antibodies by Western blotting, whereby signal intensity is directly proportional to translation levels [75]. LPS enhanced puromycin incorporation as compared to control cells in a mTOR-dependent manner, which is indicative of elevated protein synthesis in LPS-exposed macrophages (S5f and S5g Fig). Given that mTORC1 regulates mRNA translation through multiple mechanisms [39], we next examined the role of S6Ks and 4E-BPs in LPS-mediated lysosome remodelling.

First, using LY2584702, a potent pharmacological inhibitor of S6 kinases (Fig 5a; S5h Fig), we showed that S6Ks are necessary for LPS-mediated increase in protein synthesis (Fig 5a and 5b). Second, while LY2584702 treatment did not preclude LPS-induced endo-lysosome tubulation (Fig 5c and 5d), it did prevent endo-lysosome volume expansion (Fig 5e). This observation suggests that endo-lysosome tubulation and expansion can be decoupled. Similarly, inhibition of S6 kinases thwarted the LPS-mediated increase in the levels of LAMP1, LAMP2, CD63, TRPML1, ATP6V1H and ATP6V1D (Fig 5f and 5g; S3 Fig). Importantly, LY2584702 did not affect the levels of corresponding transcripts (e.g. LAMP1, ATP6V1H, ATP6V1D and B2M) in resting cells or those co-exposed with LPS (S5i Fig). Moreover, like Akti, LY2584702 did not induce autophagy, suggesting that its effects on endo-lysosome size are independent of autophagy (S5d and S5e Fig). Together, these data suggest that the mTORC1-S6 kinase axis promotes endo-lysosomal protein expression to expand the size of the endo-lysosomal network during phagocyte activation within a couple of hours activation.

**Fig 5:**
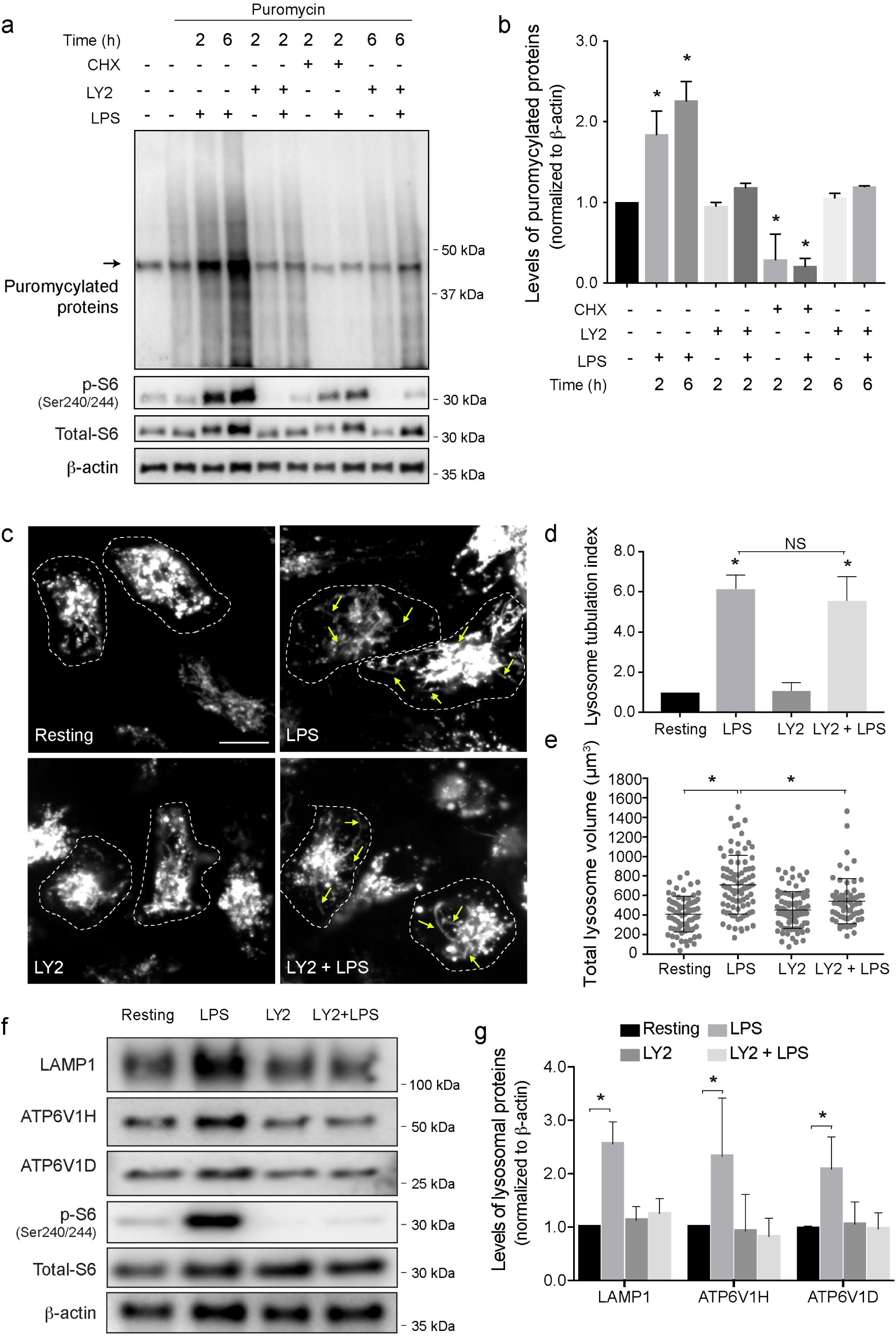
S6 kinase is required for the LPS-mediated lysosome expansion. (a) Western blot analysis of protein puromycylation in resting and activated primary macrophages. LPS increases the amount of puromycylated proteins that is blocked by p70S6K inhibitor (LY2584702) or cycloheximide. Lane 1 are control lysates from cells not exposed to puromycin. The band indicated by arrow is a non-specific band recognized by the anti-puromycin antibody. p-S6 and β-actin were used to monitor p70S6K activity and as a loading control, respectively. (b) Normalized puromycylation signal (excluding non-specific band) normalized over β-actin signal. Data is shown as the mean ± standard deviation from three independent experiments. Statistical analysis was done with an ANOVA, where * indicates conditions that are statistically distinct from control (*p<0.05). (c) Lysosomes in primary macrophages were pre-treated with LY2584702 (LY2) followed by 2 h of LPS where indicated. Images were acquired by live-cell spinning disc confocal microscopy. Scale bar = 5 µm. (d) Lysosomal tubulation was scored for each condition as shown, where a tubule was defined as longer than 4 µm in length. Tubulation index was determined by normalizing scores to resting cells. (e) Total lysosome volume in primary macrophages treated as indicated. For d and e, shown are the mean ± standard error of the mean from 30-40 cells per condition per experiment, across three independent experiments. (f) Western blot analysis of whole cell lysates from resting and activated primary macrophages with or without LY2584702. (g) Quantification of Western blots showing the levels of LAMP1 and the V-ATPase V_1_ subunits H and D, normalized to β-actin. p-S6 and total S6 blots are shown to support effectiveness of LY2584702 treatment. Shown is the mean ± standard deviation of the mean from five independent blots. For b, c and e, data was statistically analysed with ANOVA and unpaired post-hoc test (*p<0.05). For each figure with Western blots, see S1 Raw Images for original, unedited Western blots. See S5 Data for original data in Fig 5.

We next investigated the role of 4E-BPs in regulating endo-lysosome expansion following LPS. For this, we generated RAW macrophages that stably express 4E-BP1^4Ala^, a phosphorylation-deficient mutant of 4E-BP1 carrying alanine substitutions at four phosphorylation sites (Thr37, Thr46, Ser65, and Thr70), rendering it inaccessible to mTORC1 regulation [76]. This form of 4E-BP1 constitutively binds to a cap-binding protein eIF4E which prevents the assembly of the eIF4F complex thereby hindering recruitment of the ribosome to the mRNA [76]. First, relative to resting RAW counterparts, we showed that LPS augmented the endo-lysosomal volume and endo-lysosome tubulation in RAW cells expressing an empty pBabe retroviral vector (Fig 6a-6c). In contrast, LPS failed to boost the endo-lysosome volume in RAW cells that stably expressed 4E-BP1^4Ala^, though endo-lysosome tubulation still occurred (Fig 6a-6c). Second, changes in endo-lysosome volume were accompanied by corresponding alterations in endo-lysosomal protein levels (we note that RAW cells are less pronounced than primary cells in terms of tubulation, expansion and increase in protein levels). Indeed, LPS exposure caused a mild but significant increase in endo-lysosomal protein levels (LAMP1, ATP6V1H and ATP6V1D) in RAW cells expressing the empty retroviral vector, compared to resting counterparts (Fig 6d and 6e). In contrast, LPS failed to boost the levels of these proteins in RAW cells stably expressing 4E-BP1^4Ala^ (Fig. 6d and 6e). Collectively, these data suggest that the effects of mTORC1 on endo-lysosome expansion are mediated via modulation of both, S6Ks and 4E-BPs.

**Fig 6:**
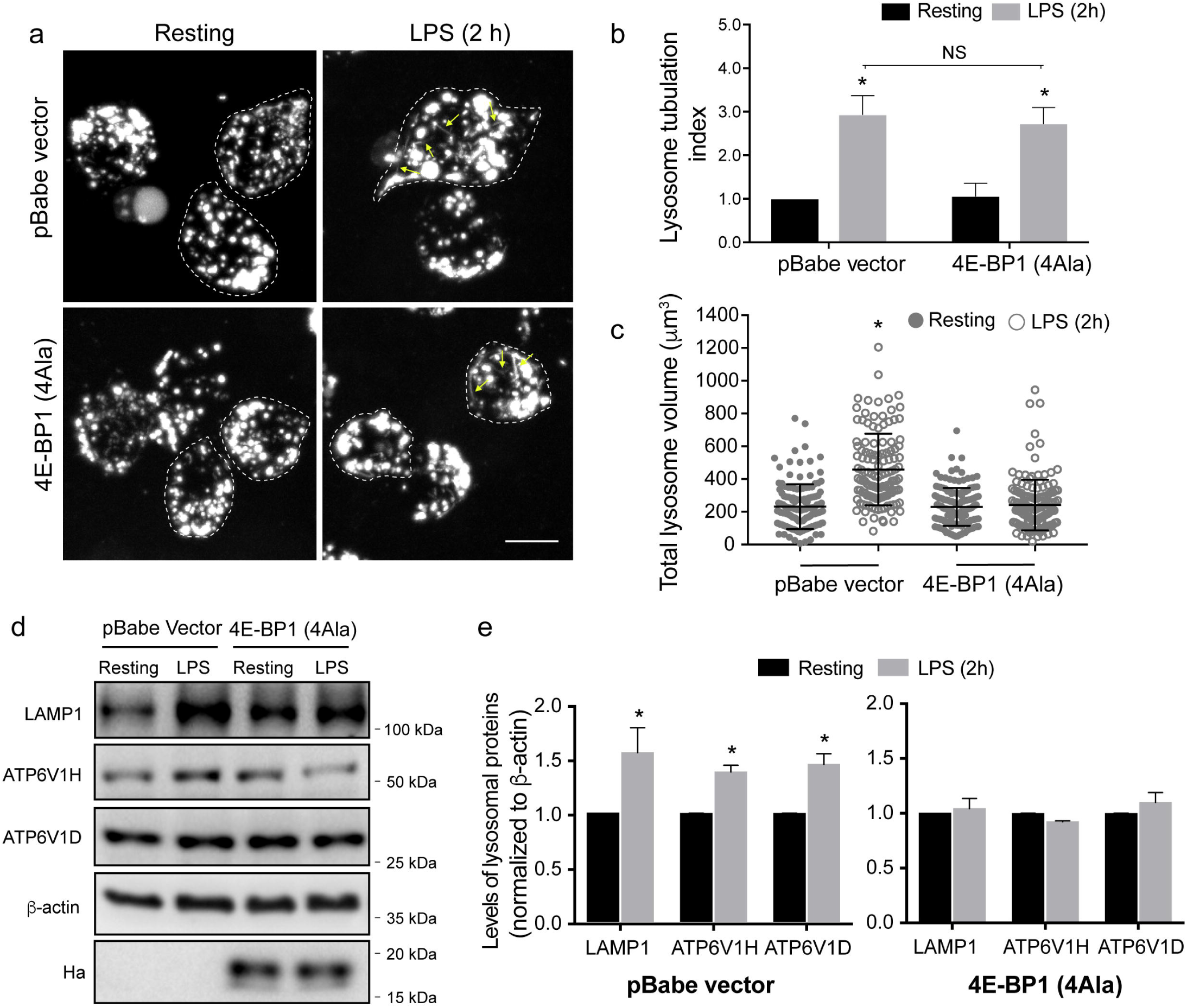
Active 4E-BP1 suppresses LPS-mediated lysosome expansion. (a) Lysosomes in resting or LPS stimulated (2 h) RAW cells stably expressing the 4E-BP1 (4Ala) phosphorylation mutant or the empty pBabe vector. Images were acquired by live-cell spinning disc confocal microscopy. Scale bar = 10 µm. (b) Lysosomal tubulation was scored for each, where a tubule was defined as longer than 4 µm in length. Tubulation index was determined by normalizing scores to resting. (c) Total lysosome volume in engineered RAW macrophages treated as indicated. For b and c, shown are the mean ± standard error of the mean from 30-40 cells per condition per experiment, across three independent experiments. (d) Western blot analysis of whole cell lysates from stable cell lines. (e) Quantification of Western blots showing the levels of LAMP1 and the V-ATPase V_1_ subunits H and D, normalized to β-actin for both cell lines. Anti-HA blot demonstrates expression of 4E-BP1^4Ala^. Shown is the mean ± standard deviation of the mean from 3 independent blots. For b, c and e, data was statistically analysed with ANOVA and and unpaired post-hoc test (*p<0.05). For each figure with Western blots, see S1 Raw Images for original, unedited Western blots. See S6 Data for original data in Fig 6.

### Polysome profiling of transcripts encoding endo-lysosomal proteins after LPS exposure

Considering that LPS increased six endo-lysosomal protein levels that we tested without increasing the corresponding mRNA abundance in primary macrophages (Figs 2 and 3; S3 Fig), we next postulated that LPS-driven mTOR activity promotes endo-lysosome expansion by enhancing translation of at least a subset of mRNAs encoding endo-lysosomal proteins. To test this hypothesis, we employed polysome profiling wherein mRNAs are separated according to the number of bound ribosomes by sedimentation through a 5-50% sucrose gradient [77]. Distribution of mRNAs encoding endo-lysosomal proteins across the gradient was measured by RT-qPCR. Due to technical limitations, these experiments were carried out using RAW macrophages. We tested polysomal distribution of mRNAs encoding LAMP1, V-ATPase subunits H and D, and cathepsin D.

Relative to the control, LPS treatment for 2 or 6 h shifted the distribution of mRNAs encoding LAMP1, and the V-ATPase subunits H and D towards the heavy polysome fractions, which is indicative of increased translational activity of these mRNAs (Fig 7a-c and S5a-c Fig). Importantly, although torin1 exerted minimal effect on the distribution of mRNAs encoding LAMP1 and V-ATPase subunits H and D in control cells (S7 Fig), it attenuated the shift of these transcripts towards heavy polysomes in LPS-treated cells (Fig 7a-c; S5a-c Fig). These findings indicate that LPS induces translation of mRNAs encoding LAMP1, and V-ATPase subunits H and D via mTOR. Notably, translational regulation of LAMP1 and V-ATPase subunits H and D is consistent with the results obtained in primary macrophages wherein LPS induced LAMP1 and V-ATPase subunits protein levels without affecting their mRNA levels or protein stability (Figs 2a and 3c; S3 Fig).

**Fig 7:**
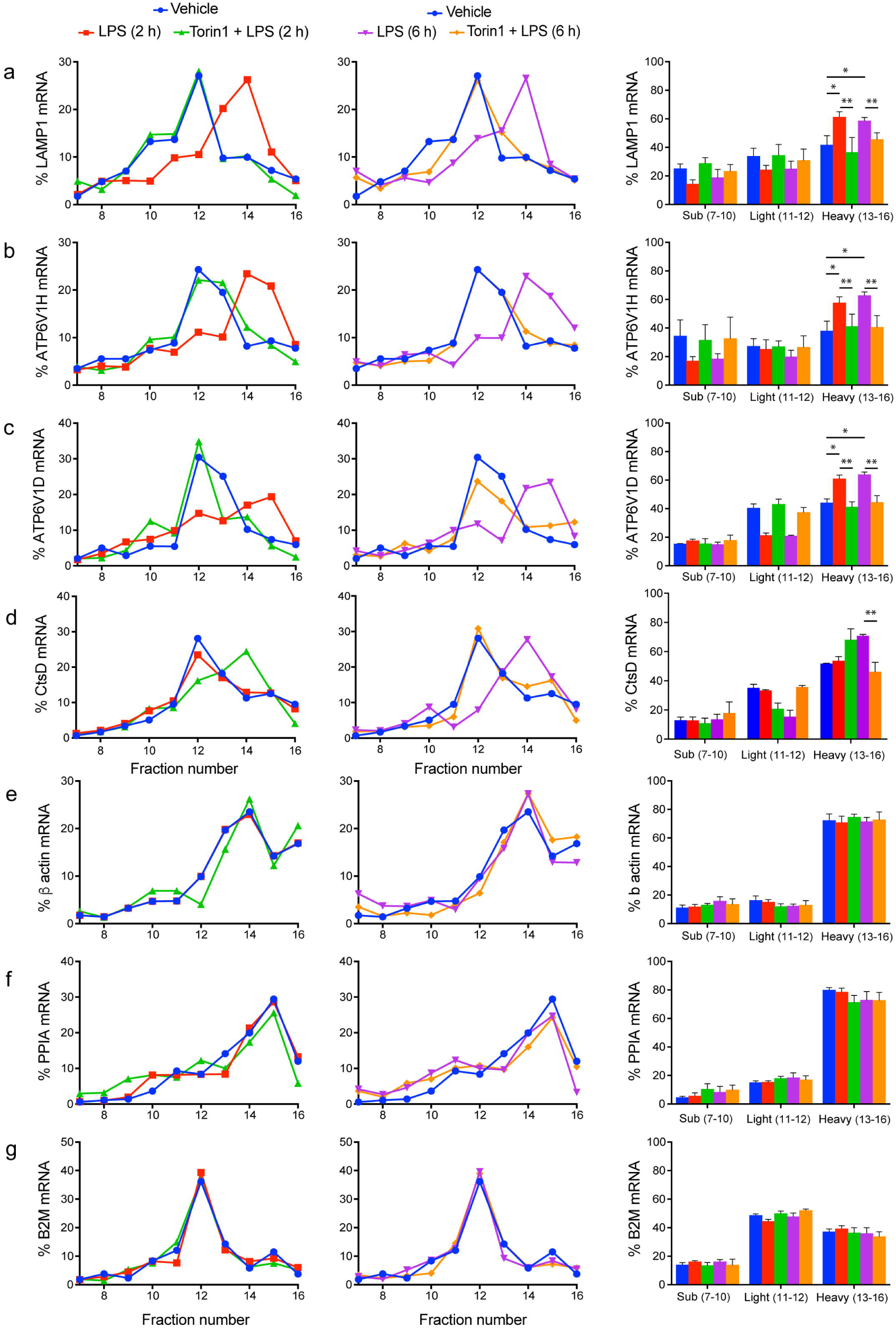
LPS increases translation of mRNAs encoding lysosomal proteins in an mTOR-dependent manner. (a-g) Percent of target mRNA (a: LAMP1, b: ATP6V1H, c: ATP6V1D, d: CtsD, e: β-actin, f: PPIA, and g: B2M) associated with each ribosome fraction in resting, LPS- or LPS/torin1-treated RAW cells. Left and middle panels show 2 h and 6 h treatments, respectively. Shown is a representative experiment from four independent experiments, each of which contained three technical replicates. Right panels: Pooled percent mRNA in subpolysomal (fractions 7-10), light polysosome (fractions 11 and 12) and heavy polysomes (fractions 13-16). Shown is the mean percent ± standard deviation from four independent experiments with each point in triplicate for each experiment and mRNA. Heavy fractions were statistically analysed by ANOVA and Tukey’s post-hoc test, where * indicates statistical difference from resting conditions, while ** indicates differences between LPS and LPS+torin1 conditions within 2 and 6 h exposure. See S7 Data for original data in Fig 6.

In comparison, mRNAs encoding cathepsin D did not shift to heavier polysome fractions after 2 h LPS treatment (Fig 7d, S5e Fig). Yet, exposure to torin1 alone or co-administration of LPS and torin1 for 2 h caused cathepsin D mRNA to shift to heavy polysomes (S7d Fig); this may be aligned with the need for increased catabolic activity during starvation conditions that repress mTORC1, stimulate autophagy, and activate TFEB [45,46,62,78]. Interestingly, and indicative that endo-lysosomes undergo different phase of remodelling during phagocyte maturation, 6 h LPS caused mRNA encoding cathepsin D to shift to heavy polysomes, while this was impaired by co-administration of LPS and torin1 for 6 h (Fig 7d, S5e Fig). These changes in translational activity of mRNAs encoding key endo-lysosomal proteins are in contrast to mRNAs encoding β-actin, peptidylpropyl isomerase A (PPIA), and β2-microglobulin (B2M), whose polysome distribution was not majorly perturbed by LPS and/or torin1 treatments (Fig 7e-g, S5f-h and S7e-g Figs). This is consistent with previous reports showing that translation of these mRNAs is insensitive to LPS and/or mTORC1 [79, 80]. Collectively, these observations suggest that translation of mRNAs encoding specific endo-lysosomal proteins is selectively modulated during macrophage activation by LPS in an mTOR-dependent manner.

### LPS induced changes of the translatome are largely mediated by mTORC1

We next employed polysome-profiling in conjunction with RNASeq to identify genome-wide changes in the transcriptome (i.e. steady-state mRNA levels which are influenced by changes in transcription and/or mRNA stability) and translatome (i.e. the pool of polysome-associated mRNAs) [77]. To allow stringent statistical analysis, we sequenced matched total transcriptomes and translatomes from three independent experiments from RAW cells untreated, exposed to LPS alone or in combination with torin1 for 6 h. Parallel sequencing of polysome-associated and corresponding total mRNA followed by anota2seq analysis enables identification of *bona fide* changes in translational efficiencies (i.e. changes in polysome-associated mRNA independent of changes in total mRNA), changes in mRNA abundance (i.e. congruent changes in total mRNA and polysome-associated mRNA) and translational buffering (i.e. changes in total mRNA that are not accompanied by alterations in polysome-associated mRNA) [81].

We first contrasted effects of LPS and combination of LPS and torin1. Consistent with the increase in mTOR activity, 6h LPS stimulation leads to ample changes at the total and polysome-associated mRNA levels as compared to resting cells (Fig 8a). LPS-induced changes in polysome-associated mRNA levels appeared to be mostly influenced by alterations in mRNA abundance; however, a subset of mRNAs exhibited perturbed translation efficiencies (Fig 8b, S1 Table). We then sought to characterize to what extent these changes in translation efficiencies are mediated by mTOR. Therefore, we visualized those subsets of mRNAs that were translationally modulated after LPS stimulation and monitored their behavior after mTOR inhibition (as in Fig 8b, but highlighted in Fig 8c and 8d). In RAW cells, torin1 lead to a near-complete reversal of the LPS-induced translatome (compare Fig 8c-d). This indicates that the effects of LPS on translational efficiency are largely mediated by mTOR. Moreover, transcripts whose polysome-association was stimulated by LPS comprised a number of TOP mRNAs which are well-established translational targets of mTOR. Thus genome-wide experiments revealed that in addition to transcriptional remodeling, 6h LPS treatment entices mTOR-dependent perturbations in the translatome.

**Fig 8:**
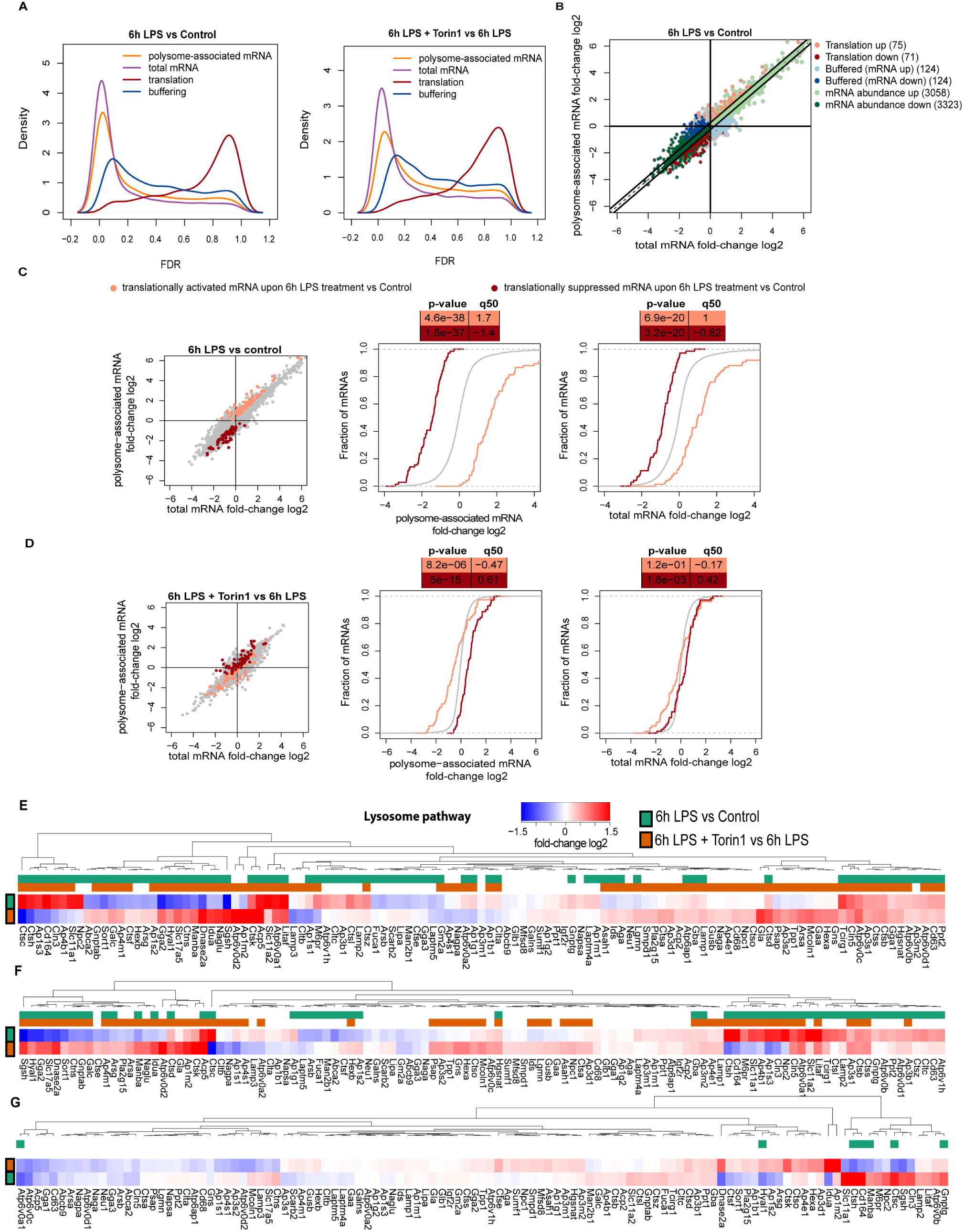
Transcriptome analysis of heavy polysomes during LPS-mediated activation of RAW cells. (a)Distributions (Kernel density estimates) of False Discovery Rates (FDRs) from comparisons of gene expression between 6h treatments with LPS to control (left); and LPS in presence or absence of torin-1. Differences in polysome-associated mRNA (orange), total mRNA (purple), translation (red) and buffering (blue) were assessed using polysome-profiling. (b) Scatterplot of polysome-associated mRNA vs total mRNA log2 fold changes (6h LPS to control). The number of transcripts exhibiting changes in translation (red), buffering (blue) and mRNA abundance (green) stratified into increased (light shade) or decreased (dark shade) expression are indicated. (c) Scatterplot of polysome-associated mRNA vs total mRNA log2 fold changes (left) together with cumulative distribution plots for polysome-associated mRNA (middle) and total mRNA (right) log2 fold changes from the comparison between 6h LPS treatment to control. Translationally activated (light red) and translationally suppressed (dark red) mRNAs are indicated together with background transcripts (i.e. not in either of sets; grey). (d) Same plots and subsets of transcripts as in **c** but using gene expression data originating from the comparison between 6h LPS treatment in presence or absence of torin-1. (e-g) Heatmap of log2 fold changes for total mRNA (e), polysome-associated mRNA (f) and changes in translation efficiencies (g) following 6h treatments with LPS relative to resting (green); and LPS in presence relative to absence of torin-1 (orange) for genes annotated to the lysosme pathway. The sidebars indicate genes with significantly changed expression in their associated analysis separately for the two comparisons (i.e. green or orange). See S1 Table and deposited data in Gene Expression Omnibus (GEO) with accession number GSE136470 for original data in Fig 8.

We then focused on mRNAs encoding lysosome-associated proteins and interrogated how their expression changed upon 6h LPS treatment relative to resting conditions (Fig 8e-g, S1 Table). This revealed subsets of mRNAs encoding lysosome proteins showing distinct patterns of regulation. For example, transcripts encoding the cholesterol transporter NPC2, the divalent metal transporter SLC11A1, and the transmembrane protein Cln3 increased, while those encoding sulfamidase (Sgsh) and Hyaluronidase-1 (Hyal1) decreased in a manner sensitive to torin1; the increase in many lysosomal mRNAs in the presence of torin-1 may be explained by TFEB activation during mTOR inhibition (Fig. 8e). Moreover, we identified 59 mRNAs that significantly changed in their association with heavy-polysomes. For instance, transcripts encoding proteins like lysosome-biogenesis receptor M6PR, SLC11A1, a second metal transporter SLC11A2, LAMP2, Cln3, CD63, NPC2, and several V-ATPase subunits were increased in their abundance in heavy polysome fractions in response to LPS (Fig 8f, S1 Table). The opposite was observed for the lysosomal enzymes like Sgsh, Hyal1, and the galactosylceramidase (Galc) (Fig 8f, S1 Table). Interestingly, a significant number of LPS-induced changes was abated or even reversed by the addition of torin1, thereby suggesting a role for mTOR in modulating association of these mRNAs with heavy polysomes (Fig 8f, S1 Table). Nevertheless, when mRNA levels in heavy polysomes were adjusted for changes in total mRNA levels to capture alterations in translation efficiency, only a few mRNAs were identified as significant but included M6PR, LAMP2, Cln3 (Fig. 8g, S1 Table). Overall, these data suggest that endo-lysosomes in addition to undergoing LPS-induced expansion over phagocyte maturation (as supported by Figs. 1 and 2), may be functionally remodelled.

To validate these findings, we performed qRT-PCR analysis of distribution of six top LPS-stimulated (NPC2, Cln3, SLC11A1, SLC11A2, M6PR, CtsC) and two top LPS-suppressed mRNAs (Sgsh, Hyul1) across unpooled gradient fractions. Consistent with the RNASeq analysis, we observed that LPS induces a shift of mRNAs encoding NPC2, Cln3, SLC11A1, SLC11A2, M6PR, and CtsC towards heavy polysome, while Sgsh and Hyal1 were depleted from heavy polysomes (S8 Fig). Both effects appeared to be sensitive to torin1 (S8 Fig). Overall, using this strategy, we observed that LPS can selectively enrich or deplete specific mRNAs encoding endo-lysosomal transcripts, which provides evidence that LPS may functionally remodel the endo-lysosomal system during phagocyte maturation in addition to its expansion (as supported by Fig. 1 and Fig. 2).

Surprisingly, the initial candidate mRNAs encoding endo-lysosomal factors that we tested (e.g. LAMP1, V-ATPase subunits H and D, and cathepsin D) were for the most part not captured by RNASeq of pooled fractions – this is despite validating a large proportion of mRNAs identified as differentially translated by RNAseq in pooled heavy polysome using qRT-PCRs across the whole gradient (e.g. NPC2, SLC11A2). This may be explained by technical issues related to polysome profiling/RNASeq. Namely, ribosome-association of mRNAs shows normal distribution with a large coefficient of variance, thereby implying that for example even if the large proportion of mRNA is associated with >3 ribosomes, a fraction of it should be associated with <3 ribosomes [82]. Moreover, based on the empirically assessed behaviour of the vast majority of cellular mRNAs, a threshold of 3 and more polysomes is set to distinguish between efficiently and not efficiently translated mRNAs [82]. Nevertheless, in cases when mRNAs shift within heavy or light polysome fractions but do not exhibit significant migration over threshold of 3 and more ribosomes, the power to detect changes in translational efficiency may be reduced. This could thus mask detecting effects on alterations of mRNA translational efficiency upon LPS stimulation of some mRNAs including those encoding LAMP1 and specific V-ATPase subunits. This is supported by targeted qRT-PCR against LAMP and V-ATPases on unpooled fractions from the same samples we used for global analysis. We again observed a shift in the abundance of mRNAs encoding these proteins to heavier polysomes upon LPS (even at 6 h) that was reduced by torin1 treatment (S9 Fig) However, the size of the response was less pronounced and was likely averaged out during pooling. As before, mRNAs encoding PPIA, actin, and B2M largely did not respond to LPS or torin1 (S9 Fig). Of note, we opted for polysome instead of ribosome profiling, as although ribosome profiling has far superior resolution inasmuch as it can determine ribosome positioning at a single nucleotide level, polysome profiling appears to perform better in determining translational efficiencies [83]. Additionally, in the case of qRT-PCR where the treatments were for 2 and 6 hours, for the RNAseq analyses only 6h time point was used.

### Antigen presentation is promoted by LPS through mTOR and S6K activity

Our results suggest that LPS can expand the endo-lysosomal system within two hours of activation in primary phagocytes via an mTOR-dependent alterations in translation of key endo-lysosomal transcripts. We next sought to investigate the functional implication of this LPS-mediated escalation in translation and endo-lysosomal volume. Given that antigen processing and loading occurs within the endo-lysosomal of DCs, we postulated that LPS-mediated expansion of endo-lysosome volume and holding capacity may enhance antigen presentation in BMDCs. To test this, we used BMDCs from C57Bl/6 and C3H/He mice that respectively carry MHC-II haplotypes I-A^b^ and I-A^k^. I-A^b^ and I-A^k^ expressing BMDCs were then respectively fed antigens, the peptide Eα^52-68^ and full-length Hen Egg Lysozyme (HEL), for 4 and 6 hours in the presence or absence of LPS. Fixed, but unpermeabilized BMDCs were then stained with Aw3.18 monoclonal antibodies to detect surface delivery of I-A^k^:: HEL^48-62^ complexes [84] – this antibody could not detect MHC-II:peptide complex after permeabilization (not shown). On the other hand, we could detect total I-A^b^::Eα^52-68^ [85] (internal and surface level) complex formation by staining fixed and saponin-permeabilized BMDCs with the monoclonal Y-Ae antibody [85]. Importantly, treatment with LPS stimulated formation and/or delivery of the MHC-II::peptide complexes even at 4 h and more potently at 6 h (Fig 9a-c and S10 Fig). When cells were not given antigens, the signal was reduced to background, demonstrating that the fluorescence signal was dependent on MHC-II::antigen complex formation (Fig 9c, S10 Fig.). We then inquired if antigen presentation was dependent on mTOR and S6K activities by co-treating cells with torin1 and LY2584702, respectively. Remarkably, these inhibitors reduced antigen presentation of both Eα^52-68^ and HEL^48-62^ in unstimulated and LPS-treated cells (Fig 9a-c and S10 Fig.). These data indicate that altered translation controlled by the mTORC1-S6K axis is necessary for efficient antigen presentation by BMDCs. Consistent with these data, we observed that LPS boosted total MHC-II levels in BMDCs and that this was prevented by treatment with torin1 and LY2584702 (Fig 9d and 9e). These observations suggest that enhanced translation driven by mTORC1 and S6Ks underpins, at least in part, boosting MHC-II levels in BMDCS.

**Fig 9:**
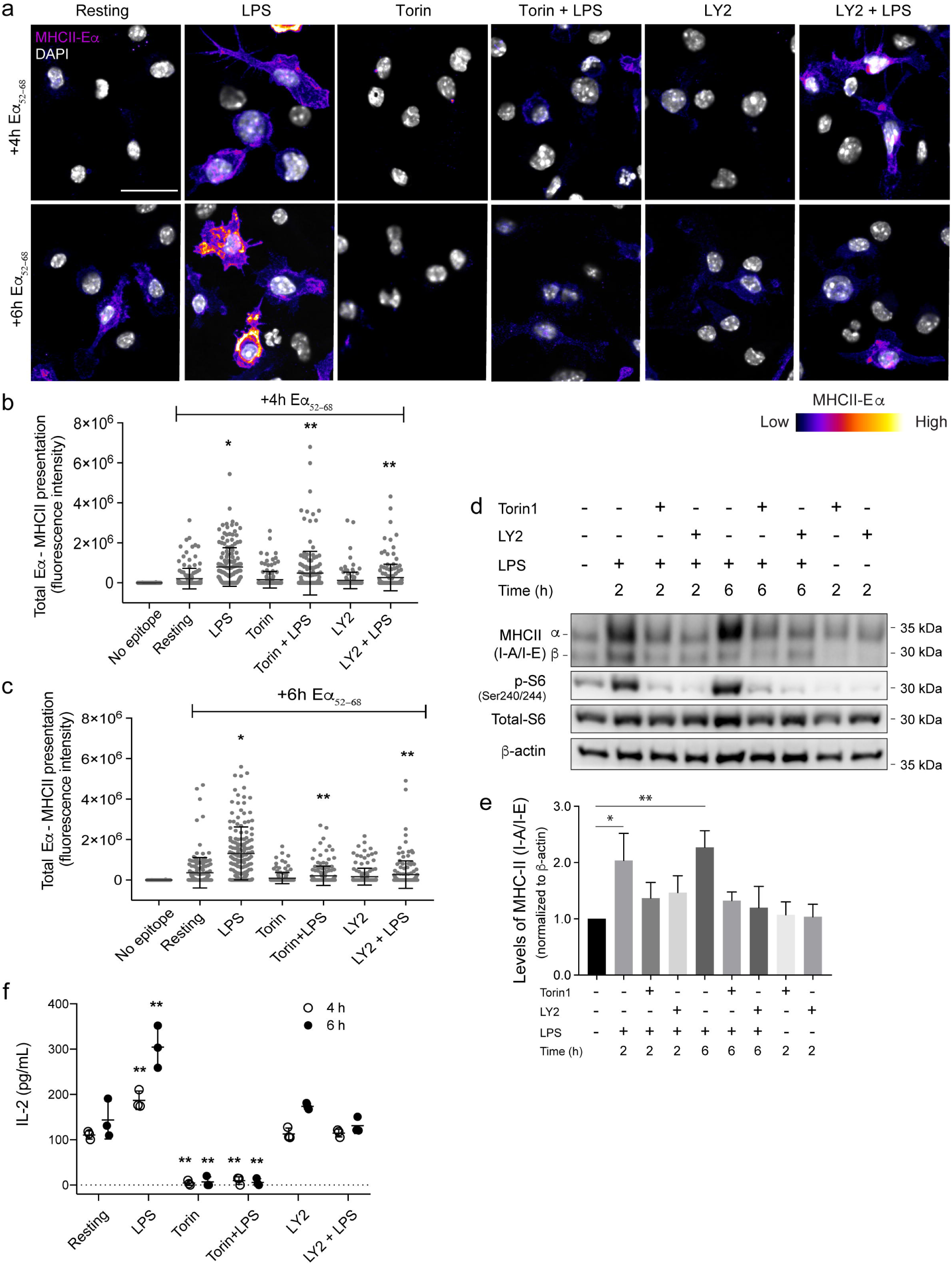
mTOR and S6 kinase control Eα^52–68^ peptide presentation in activated BMDCs. BMDCs were incubated with Eα^52–68^ peptide for 4 or 6 hours in the presence or absence LPS with or without torin1 and LY2584702. Cells were then fixed and stained with Y-Ae antibodies to detect I-A^b^::Eα^52–68^ complex formation, and DAPI to stain nuclei. (a) I-A^b^::Eα^52–68^ complexes (displayed in pseudo-colour) and DAPI (grayscale) are shown for BMDCs treated as indicated. (b, c) Anti-I-A^b^::Eα^52–68^ antibody signal was quantified by fluorescence intensity associated with each cell. Shown is the mean of the total fluorescence intensity of I-A^b^::Eα^52–68^ complexes ± standard deviation from three experiments, where 50-100 cells were quantified for each. Data was analysed using ANOVA, whereby * indicates a difference compared to unstimulated BMDCs exposed to Eα^52–68^ and ** indicates a difference compared to LPS-stimulated BMDCs fed Eα^52–68^ (p<0.05). Scale bar = 30 µm. Color scale: 0 – 2500 (low-high). S10 Fig show similar data for HEL presentation. (d) BMDCs were fed Eα^52–68^ peptide for 4 or 6 hours in the presence or absence LPS with or without torin1 and LY2584702. Following mild fixation, APCs were co-incubated with T-cells as described in methods, to measure I-A^b^::Eα^52–68^ complex induced T-cell activation. T-cell secreted Il-2 was measured using an ELISA system. All data were analysed using ANOVA, whereby ** indicates a difference compared to unstimulated BMDCs exposed to Eα^52–68^ (p<0.05). (e) Western blot analysis of whole cell lysates from APCs. p-S6 and β-actin were used to monitor mTOR-p70S6K signaling axis activity and as a loading control, respectively. (f) Quantification of Western blots showing the levels of MHC-II (I-A/I-E) normalized over β-actin signal. Data is shown as the mean ± standard deviation from four independent experiments. Statistical analysis was done with an ANOVA, where * and ** indicates a significant difference of 2 hour and 6 hour conditions respectively, from resting cells (p<0.05). For each figure with Western blots, see S1 Raw Images for original, unedited Western blots. See S8 Data for original data in Fig 9.

To determine whether LPS-mediated stimulation of mTORC1/S6K axis was needed for antigen-mediated activation of T cells, we measured IL-2 release by T cells to assess activation. For this assay, we used the T cell hybridoma clone 1H3.1, which recognizes I-A^b^::Eα^52-68^ [86]. BMDCs were fed Eα^52-68^ and then either treated with vehicle, or exposed to torin1 or LY2584702 alone, or to LPS with and without drugs. We also included BMDCs without any antigen feeding. After 6 h of antigen processing and presentation, BMDCs were fixed and co-incubated with 1H3.1 T cell hybridomas for 48 h. We then employed ELISA to quantify IL-2 released by T cells. T cells released IL-2 at greater levels when they were exposed to BMDCs treated with both peptide and LPS than when they were exposed to BMDCs fed with only peptide alone (Fig 9f). This activation was dependent on Eα^52-68^ as very little IL-2 was discharged by T cells exposed to BMDCs that were not fed peptide (dashed line). In comparison, IL-2 secretion was nearly at baseline when T cells were co-incubated with APCs treated with LPS and torin1, demonstrating that mTORC1 is critical for antigen presentation by BMDCs. Importantly, BMDCs pre-treated with LY2584702 caused T cells to secrete IL-2 at levels similar to BMDCs that internalized only peptide (Fig 9f). This result demonstrates that enhanced protein synthesis driven by S6Ks is critical for APCs to stimulate antigen presentation and T cell activation. Though we acknowledge that increased protein synthesis may play additional roles in phagocytes that lead to enhanced antigen presentation, collectively our data suggest that LPS-driven protein synthesis and lysosome expansion are essential to boost antigen retention and ultimately presentation.

## Discussion

Macrophages and dendritic cells are plastic cells inasmuch as they can alter their metabolic and gene expression profiles to adopt a range of alternative states, which exert either inflammatory or anti-inflammatory functions. Significant attention has been given to how macrophages and dendritic cells alter their metabolism and expression of cytokines, chemokines and other microbicidal agents [8–11]. Remodelling the expression level of these factors occurs both at the transcription and translation level [49,70,87,88]. However, remarkably less is understood regarding the mechanisms that underpin changes to the endomembrane system during activation of macrophages and dendritic cells. Notable examples of changes to the endomembrane system include reduced degradative capacity of endosome and lysosomes to help conserve antigens in dendritic cells, delayed phagosome maturation in interferon-γ-treated macrophages, and a morphological reorganization of the endo-lysosome system in both cell types, shifting from a large collection of vesicular organelles into a tubular network of endo-lysosomes [13,21,22,89]. Tubular endo-lysosomes are associated with increased pinocytic retention, exchange of phagosomal content within the endo-lysosomal system, and delivery of MHC-II-peptide for presentation, though how this occurs remains unknown [17,20,21,24,90]. Herein, we show that phagocytes also expand the endo-lysosome system and its holding capacity within two hours of activation. We provide evidence that this expansion relies on altered translation activity driven by the mTORC1-S6K-4E-BP axis to boost expression of several important endo-lysosomal proteins like LAMP1, LAMP2, TRPML1, and V-ATPase subunits. In turn, this seems to enhance antigen retention, leading to efficient antigen presentation.

### Expansion and reorganization of the endo-lysosome system in activated phagocytes

Here, we disclose that LPS-activated macrophages and dendritic cells remodel their endo-lysosome system into an expanded tubular network with augmented holding capacity. This conclusion is supported by several observations. First, imaging volumetric analysis revealed that dyes preloaded into endo-lysosomes occupy a greater volume post-LPS activation using both live- and fixed-cell imaging. This increase was not due to an artifact caused by imaging geometrically distinct objects since collapse of tubular endo-lysosomes during standard fixation with 4% PFA or the use of super-resolution imaging still captured a larger volume in LPS-treated phagocytes than their resting counterparts. Second, there was a significant increase in the expression level of major endo-lysosomal proteins including LAMP1, LAMP2, CD63, TRPML1, and at least two V-ATPase subunits that was blunted by cycloheximide treatment or when up-regulation of translation was prevented by inhibiting Atk, mTOR, S6Ks, or 4E-BPs. In dendritic cells, this included an enhanced expression of MHC-II subunits. Third, activated macrophages could hold a larger amount of fluid-phase relative to resting counterparts, suggesting a larger endo-lysosomal compartment to store pinocytic cargo. Thus, overall, activated phagocytes not only undertake morphological reorganization of endo-lysosomes but also expand this organelle network. The increase in endo-lysosome volume and holding capacity is consistent with work by Swanson *et al.* done in the 1980s showing that phorbol ester-activated macrophages retain fluid-phase more effectively than resting macrophages [20, 61]. Thus, we argue that activated phagocytes expand their endo-lysosomal system.

### Functional implications of endo-lysosome expansion and reorganization

Functionally, an expanded endo-lysosome volume may help phagocytes engulf more material and/or process extracellular particulates and soluble cargo more efficiently, as supported by our observation that macrophages can accumulate larger amounts of pinocytic tracers upon LPS-mediated activation. This expansion of the endo-lysosome system should benefit both macrophages and dendritic cells upon activation. While mature dendritic cells have been reported to have reduced endocytosis [56, 57], we show here that dendritic cells exhibit significant pinocytic activity for at least 8 h post-activation, providing an avenue to internalize and accumulate antigenic material earlier during maturation. This is also consistent with recent reports revealing that dendritic cells are still able to internalize significant amounts of extracellular content [58, 59]. In agreement with this concept, we showed here that LPS stimulation of BMDCs increased presentation of two distinct antigenic peptides (HEL^48-61^ and Eα^52-68^) and activation of cognate T cell lines as early as 4-6 h of antigen uptake. Importantly, efficient antigen presentation was dependent on mTOR and S6K activities expressed in dendritic cells (as opposed to T cells), suggesting that up-regulated translation coupled to endo-lysosome expansion helps drive antigen presentation. Culminating these observations, we showed that mTOR and S6Ks are important for LPS-activated BMDCs to prime cognate T cells.

Additional processes may facilitate antigen presentation in phagocytes including LPS-mediated alteration of lysosomal properties like pH, redox state, and degradative capacity. Indeed, our observations suggest that the endo-lysosomal system does not uniformly expand but may undergo a reorganization in its composition and function, and that this may itself change over time during phagocyte maturation. For example, we observed that the levels of cathepsin D did not change during 2 h LPS activation of primary macrophages, unlike other proteins we examined (Fig 2 and S3 Fig). Moreover, transcripts encoding cathepsin D did not accumulate in heavy polysomes during 2 h LPS in RAW cells, but then accumulated after 6 h of LPS (Fig 7). This was corroborated by the genome-wide landscape of mRNAs associated with heavy polysomes in response to 6 h LPS treatment (Fig 8 and S8 Fig). For example, our observation that SLC11A1 and SLC11A2 are enriched in heavy polysomes in response to LPS is tantalizing since these proteins sequester divalent Fe^2+^ from microbes as an anti-microbicidal activity [91]. Thus, we propose that endo-lysosomes not only expand, but undergo dynamic functional reorganization over phagocyte maturation. However, we note the caveat that the genome-wide polysome profile analysis was completed using RAW cells, a transformed cell line that grows rapidly and may thus have distinct properties than non-proliferative primary phagocytes. Thus, it will be important to develop and employ transcriptome, polysome profiling, and proteomic analyses using primary phagocytes to better assess phagocyte maturation and adaptation of their endomembrane system.

### Acute (2 h) endo-lysosome expansion is not likely driven by transcription

Our data suggest that acute (2 h) upward scaling of the endo-lysosome system in response to LPS is not associated with transcription upregulation of genes encoding endo-lysosomal proteins. First, while TFEB and TFE3 can scale up lysosomal activity in response to various stresses [14,66,92,93], their kinetics of activation by LPS did not mirror those of endo-lysosome enlargement; endo-lysosome expansion was achieved within 2 h (Figs 1 and 2; S3 Fig), while nuclear entry of TFEB/TFE3 required 6 h post-LPS exposure (Fig 3), consistent with past observations [66]. This delayed activation suggests that TFEB and TFE3 are stimulated indirectly by LPS exposure. Second, deletion of TFEB and/or TFE3 did not impair tubulation or endo-lysosome expansion, at least in RAW cells (Fig 3). Third, we did not observe induction of mRNA levels encoding six major lysosomal proteins 2 h post-LPS exposure in primary macrophages, suggesting that enhanced transcription was not responsible for increased protein levels of LAMP1, LAMP2, CD63, TRPML1, and V-ATPase subunits within this time frame (Figs 1 and 3c; S3 Fig). Nevertheless, there are two key caveats in our study. For one, the above conclusions are limited to a select few mRNAs in primary macrophages. Transcriptome analysis of wild-type and *tfeb*^−/−^ primary macrophages will need to be completed to better understand the contributions of TFEB to endo-lysosome remodelling in response to LPS. Second, we limit our conclusions to early lysosome remodelling, defined here as 2 h of LPS. Transcriptional processes may become more pronounced at latter times to remodel endo-lysosomes, as suggested by transcriptome analysis after 6 h of LPS (Fig 8).

### Acute endo-lysosome expansion is driven by translational up-regulation

Phagocyte activation expands the endo-lysosome system within two hours of LPS exposure. This enlargement is driven by *de novo* protein synthesis as indicated by the cycloheximide-mediated block of the endo-lysosome expansion. Importantly, we observed that mTOR-dependent translational mechanisms play a key role in LPS-mediated endo-lysosome expansion. First, LPS activates mTORC1, as supported by increased phosphorylation of S6K and 4E-BP1, and enhanced global protein synthesis; importantly, inhibition of mTOR abrogated endo-lysosome expansion. Second, while the translation machinery is governed by a plethora of mechanisms, including mTOR-independent pathways, our data revealed that S6Ks and 4E-BPs play a major role in governing endo-lysosome size expansion in response to stress. mTORC1-mediated inhibition of 4E-BPs releases the translation brake imposed on the translation initiation factor eIF4E [39, 94]. On the other hand, the role of S6Ks in this process is potentially more complex given its numerous targets that modulate translation, including the ribosomal protein rpS6, the translation initiator factor eIF4B, PDCD4 which governs the function of eIF4A, eEF2K which governs elongation rates and SKAR that may promote mRNA splicing and maturation [39,95,96]. It is tempting to propose that mTORC1-regulated mRNA translation may broadly serve to rapidly scale the activity and size of other organelles in response to various signals that regulate cell differentiation, metabolic re-wiring, and stress resolution. Consistent with this idea, inhibition of TSC1/2, an inhibitor of mTORC1, or overexpression of S6Ks increased the size and length of cilia in *Chlamydomonas reinhardtii* and zebrafish [97].

In addition, it is well accepted that mTOR can selectively modulate translation of specific mRNA subsets. Of these, the best characterized mRNAs are those carrying a 5’ terminal oligopyrimidine tract (5’TOP) which renders translation of corresponding transcripts mTOR-sensitive [94,98–100]. Notably, a study of human transcripts encoding 5’-TOP sequences showed an enrichment for transcripts encoding lysosomal proteins [101]. Using transcripts encoding several lysosomal proteins as proxies for endo-lysosome remodelling, we showed that early LPS-mTORC1 signaling increases translation of specific mRNAs (LAMP1, V-ATPases, NPC2, Cln3, SLC11a2, and CtsC), while not affecting cathepsin D, actin, PPIA, and B2M. While the RefSeq data base suggest that these murine mRNAs do not contain classical 5’TOP sequences, several recent studies show that a significant number of mTOR-sensitive mRNAs lack the 5’TOP motif [82, 83]. Moreover, the 5’UTRs annotated in the RefSeq database may not be those expressed [82]. We plan to establish translational assays for primary phagocytes to identify mRNA elements which guide protein synthesis during expansion of the endo-lysosomal system [82,102,103].

### A model for endo-lysosome remodelling in activated phagocytes

Acute LPS-mediated phagocyte stimulation causes extensive endo-lysosome reorganization, featuring *i)* the previously recognized morphological transformation into a tubular network and an expansion of the endo-lysosomal network, that we discovered here. We propose a model whereby mTOR independently coordinates distinct, parallel pathways to modulate endo-lysosome expansion, tubulation and possibly secretion to mediate antigen presentation (S11 Fig). Specifically, we envision that mTORC1-S6Ks-4EBPs catalyse endo-lysosome expansion through increased protein synthesis, possibly using selective translation of specific mRNAs. In comparison, tubulation and secretion of antigen-containing compartments may be driven by the mTORC1-Arl8b-kinesin pathway.

Supporting this model: *i)* BMDCs-treated with torin1 were entirely blocked for antigen presentation and T cell activation, yet those-treated with S6K inhibitors exhibited antigen presentation and T cell activation comparable to BMDCs fed antigens without LPS (Fig 9). Additionally, while torin1 potently arrests both expansion and tubulation, S6Ks and 4EBPs manipulation prevented expansion but not tubulation. This implies that mTOR plays additional roles in lysosome remodelling, while S6Ks-4EBPs drive expansion; *ii)* endo-lysosomes tubules grow towards the cell periphery and form transport intermediates that deliver antigens to the cell surface [21, 90]. It is likely that this process is controlled by the Arl8b GTPase, which couples lysosomes to the kinesin motor proteins [104]. For one, Arl8b is required for lysosome tubulation and antigen presentation [17, 105]. Additionally, LPS enhances the levels of Arl8b levels on lysosomes in an mTOR-dependent manner [17]. Collectively, we envision that LPS-driven activation of mTOR modulates several pathways to aid in endo-lysosome remodelling, culminating in enhanced antigen uptake, processing and presentation. Of course, additional processes may be at play including mTOR-modulation of V-ATPase and lysosome positioning machinery that are linked to phagocyte maturation [9, 106]. This model and the contributions made by endo-lysosome tubulation, expansion, and luminal remodeling towards antigen presentation and infection resolution will need to be assessed in future studies.

There are of course limitations to our study. First, while mTOR and S6K inhibition blocks lysosome expansion, antigen presentation, and T cell activation, and there is a strong dependence of antigen presentation on the endo-lysosomal system, the link between endo-lysosomal expansion and antigen presentation is currently correlative. It remains possible that translational activity may regulate additional pathways independently of the endo-lysosomal system that impact antigen presentation. Second, our observations suggest that expansion is not uniformly applicable to all endo-lysosomal components, suggesting that the expanded endo-lysosomal system is functionally remodelled. In this study, we did not functionally examine this, nor how this may change over the temporal scale of phagocyte maturation. Finally, our polysome profiling was done in RAW cells but with new methodologies this could be done in primary phagocytes in future studies, following how this changes over time as phagocytes mature. Nevertheless, despite these limitations, our work collectively demonstrates that activated phagocytes reorganize their endo-lysosomal system by expanding and forming a tubular network. This amplifies the endo-lysosome holding capacity of phagocytes, augmenting their ability to retain more extracellular cargo, likely contributing to enhanced antigen presentation. We demonstrate that this process is rapid, occurring within 2 hours of activation, and proceeds via enhanced and possibly selective translation of mRNAs encoding endo-lysosomal proteins, governed by mTOR, S6K and 4E-BPs. Collectively, we propose that mTORC1 and the regulated translation machinery is an important mechanism employed by cells to scale and adapt the size and volume of organelles in response to stress signals.

## Materials and Methods

### Ethics Statement

All animals were used following institutional ethics requirements under the animal user permit ACC696 and ACC907 approved by the Ryerson University Animal Care Committee, which is certified by the Canadian Council of Animal Care and the Ontario Ministry of Agriculture, Food, and Rural Affairs. Briefly, mice were anesthesized with 5% isoflurane administered by inhalation, followed by cervical dislocation before limb bone dissection to obtain bone marrow. No experiments were performed on live-animals.

### Cell lines and primary cells

Murine RAW macrophage cell lines carrying CRISPR-mediated deletion of TFEB, TFE3 or both were a kind donation from Dr. Rosa Puertollano, NIH, and were previously described [66]. These cells and the wild-type RAW264.7 (TIB-71 from ATCC, Manassas, Virginia) were grown in DMEM supplemented with 5% heat-inactivated fetal bovine serum (Wisent, St. Bruno, Canada) at 37°C with 5% CO_2_. BMDCs and bone marrow-derived macrophages (BMDMs) were harvested from wild-type 7-9-week-old female C57BL/6J mice or C3H/HeN mice (Charles River Canada, Montreal, QC) as previously described with minor modifications [107, 108]. Briefly, bone marrow was isolated from femurs and tibias through perfusion with phosphate-buffered saline (PBS) using a 27G syringe. Red blood cells were lysed using a hypoosmotic treatment. For BMDCs, cells were plated at 2 × 10^6^/well in 4 ml of DMEM supplemented with 10% fetal bovine serum, 55 µM β-mercaptoethanol, 10 ng/ml recombinant mouse granulocyte-macrophage colony-stimulating factor (PeproTech, Rocky Hill, NJ), and penicillin/streptomycin antibiotics (Wisent). Media was changed every 2 days by replacing half of the medium with fresh medium. For BMDMs, cells were plated according to experimental requirements in DMEM supplemented with 10% fetal bovine serum, 20 ng/ml recombinant mouse macrophage colony-stimulating factor (Gibco, Burlington, ON), and penicillin/streptomycin antibiotics. Media was changed every 2 days. Experiments were conducted on days 7–9.

### RAW 4EBP^4Ala^ 923 stable cell line production

We generated RAW cells stably expressing the HA-4E-BP1 (4Ala) phosphorylation mutant or the corresponding empty pBABE vector as previously described [76], with minor modifications. Briefly, pBABE constructs were transiently transfected into the 293Phoenix-AMPHO packaging cell line using Lipofectamine 2000 (ThermoFisher), as per manufacturer’s guidelines. Following 48 h, the viral titer was harvested and passed through a 0.45 μm filter. The virus-containing medium was then used to infect RAWs in the presence of 8 μg/mL polybrene (Sigma-Aldrich) for 24 h. Infection was repeated twice more. Twenty-four hours after the final infection, the medium was supplemented with 3 μg/mL puromycin (Sigma-Aldrich) and cells were selected for 1 week then harvested.

### Rate, retention and accumulation of pinocytic probes

To measure pinocytosis rate or the accumulation of pinocytic cargo, BMDMs and RAW macrophages were pulsed with 1 mg/mL Lucifer yellow (ThermoFisher Scientific, Burlington, ON) for the indicated time with and without LPS, or after 2 h of LPS pre-stimulation. For pinocytic retention, BMDMs and RAW macrophages were maintained in resting conditions or stimulated with LPS for 2 h, followed by a 30-min pulse with 1 mg/ml Lucifer yellow. Cells were then washed 3x with PBS, and fresh medium was added for the indicated chase periods. In all cases, cells were then washed in PBS, fixed with 4% PFA for 15 minutes and washed in PBS. The amount of Lucifer yellow in RAW macrophages was then quantified using LSRFortessa X-20 cell flow cytometer (BD Biosciences, Mississauga, ON) in 10,000 cells per condition per experiment. Flow cytometry analysis was performed using FCS Express 5 (De Novo Software, Los Angeles, CA). For primary macrophages, Lucifer yellow-labelled cells were visualized using ImageXpress Micro Widefield High Content Screening System (Molecular Devices, Sunnyvale, CA) by where 3×4 quadrants per well were acquired, and the level of probe was analysed using MetaXpress 6 (Molecular Devices). To analyze the pinocytic capacity of BMDCs following activation, cells were pre-stimulated with LPS for the indicated periods, followed by co-incubation with 50 µg/mL of fluorescent dextran in the remaining 30 min of the treatment. Cells were then washed 3x with PBS and fixed with 4% PFA for 15 minutes. Afterwards, dextran fluorescence was imaged by confocal microscopy and quantified with Volocity 6.3.0 image analysis software (PerkinElmer, Bolton, ON) by integrating intensity of dextran.

### Endo-lysosome labelling and tubulation

For endo-lysosome labeling, cells were pulsed with 50-100 µg/ml Alexa^546^-conjugated dextran (ThermoFisher) for 0.5-1 h, followed by 3x wash with PBS and incubated with fresh medium for at least 1 h. To induce endo-lysosome remodeling, BMDMs and BMDCs were exposed to 100 ng/mL LPS from *Salmonella enterica* serotype minnesota Re 595 (Sigma-Aldrich, Oakville, ON), while RAW macrophages were incubated with 500 ng/mL for 2 hours (unless otherwise stated). As noted earlier, we use the term “endo-lysosomes” to reflect that this labelling method likely stains the spectrum between late endosomes, lysosomes, and their hybrids, endolysosomes. For pharmacological inhibition, cells were pre-incubated for 15-20 minutes with 100 nM torin1 (Tocris Bioscience, Minneapolis, MN), 10 µM cycloheximide (Bio-Shop), 1 µM LY2584702 (Selleck Chemicals, Houston, TX) or equivalent volume of vehicle. Cells were then imaged live (unless otherwise indicated) in complete medium. Lysosome were scored as tubules if their length was greater than 4 μm.

### Antigen presentation assays

For presentation of Eα_52–68_ peptide, C57BL/6 mice with I-A^b^ background were used to isolate monocytes for BMDC differentiation, and C3H/HeN mice with I-A^k^ background (Charles River Canada, Kingston, ON) were used for presentation of Hen-egg lysozyme (HEL). Immature BMDCs were plated on Poly-D-lysine coated glass coverslips prior to incubation with model antigens. On day 7 of differentiation, dendritic cells were incubated with 2 mg/mL of HEL (Sigma-Aldrich) or 60 µM Eα_52–68_ peptide (MyBioSource, San Diego, CA) in the presence or absence of inhibitors and/or LPS, for the time points indicated.

For surface detection of I-A^k^::HEL^46–62^ complexes, Aw3.18.14 mAb was isolated from the supernatant of hybridoma B-lymphocytes (ATCC, Manassas, VA). Briefly, cells were washed with ice-cold PBS 3 times, and incubated in ice-cold Aw3.18.14 for 30 minutes, then washed with PBS and fixed in 4% PFA for 30 minutes on ice. Following, cells were incubated in Dylight-conjugated donkey polyclonal antibody against mouse (1:500; Bethyl), in standard blocking buffer for 1 hr. For presentation of I-A^b^::Eα_52–68_ complexes, cells were washed 3 times with PBS and fixed in 4% PFA for 20 mins at RT. After, cells were permeabilized in 0.1% saponin in standard blocking buffer for 1 h. Following, cells were incubated in 1:75 mAb YAe (Santa Cruz Biotechnology, Dallas, Tx) in blocking buffer for 1 hour at RT, washed with PBS and then incubated Dylight-conjugated donkey polyclonal antibodies against mouse (1:500; Bethyl), in standard blocking buffer for 1 hr. Antigen presentation of both I-A^k^::HEL^46–61^ and I-A^b^::Eα^52–68^ complexes was visualized using confocal microscopy.

### Immunofluorescence and Fluorescence Microscopy

To fix and preserve lysosome tubules in RAW cells, cells were incubated with 0.45% (v/v) glutaraldehyde and 0.5% PFA (v/v) in PBS for 15 minutes at room temperature. Cells were then washed with PBS 4x, followed by incubation with 1 mg/mL ice-cold sodium borohydride (Sigma-Aldrich) for 5 min 3x to abate unreacted glutaraldehyde and quench its autofluorescence.

To visualize endogenous TFEB and TFE3, cells were fixed using 4% PFA for 15 min following treatment conditions. Cells were then treated with 100 mM glycine in PBS to quench PFA, then in permeabilization buffer (0.2% Triton-X, 2% BSA in PBS) for 10 min and then blocked for 1 h in 2% BSA. Cells were incubated with rabbit anti-TFEB (1:200; Bethyl Laboratories, Montgomery, TX) or rabbit anti-TFE3 (1:500; Sigma-Aldrich) antibodies for 1 h, followed by Dylight-conjugated donkey polyclonal antibodies against rabbit (1:500; Bethyl) for 1 h. Nuclei were counter stained with 0.4 μg/mL of DAPI. For staining LAMP1, dextran-loaded cells were fixed in 0.45% (v/v) glutaraldehyde and 0.5% PFA (v/v) in PBS for 15 minutes at room temperature. Cells were washed with PBS 3x and quenched in 25mM glycine for 15 mins at room temperature. Cells were permeabilized in ice-cold methanol for 3 minutes and blocked in 2% BSA for 1 h. Cells were then incubated in primary rat anti-LAMP1 (1:100; Developmental Studies Hybridoma Bank) and secondary Dylight-conjugated donkey polyclonal antibodies against rat (1:500; Bethyl) for 1 h each. Cells were then mounted on a slide using DAKO mounting medium.

Live-cell imaging was done at 5% CO_2_ and 37 °C using environmental control chambers. Live-cell and fixed-cell imaging was done with a Quorum Diskovery spinning disc confocal microscope system equipped with a Leica DMi8 microscope connected to an Andor Zyla 4.2 Megapixel sCMOS or an iXON 897 EMCCD camera, and controlled by Quorum Wave FX powered by MetaMorph software (Quorum Technologies, Guelph, ON). We also used an Olympus IX81 inverted microscope equipped with a Hamamatsu C9100-13 EMCCD camera and controlled with Volocity 6.3.0 (PerkinElmer). For super-resolution imaging, we employed the Zeiss Elyra PS1 imaging system equipped with an Axio Observer Z1 microscope fitted with the Andor iXon3 885 detector for structure illumination microscopy (SIM) and powered by Zeiss Zen 2012 software (Zeiss Microscopy, Jena, Germany). Super-resolution image acquisition was acquired by grating for 3 rotations and 5 phases. All SIM reconstructed imaging was done using default settings for image reconstruction; to avoid artifact formation, only images with peak/mean ratios above 20 and noise filter less then −4 were accepted. After reconstruction, Volocity 6.3.0 (PerkinElmer) image analysis software was used. All microscopes were equipped with standard filters appropriate to fluorophores employed in this study, optics and stage automation.

### Image analysis and volumetrics

The nuclear-to-cytosolic ratio of TFEB and TFE3 was estimated as the ratio of the mean fluorescence intensity in the nucleus over the mean intensity in the cytosol after background correction using ImageJ (v. 1.47 bundled with 64-bit Java). For Lamp1 and dextran colocalization, we used Mander’s colocalization analysis to measure the degree of dextran colocalizing in LAMP1 structures, using the JACoP plugin in ImageJ after applying background subtraction. For volumetric analysis, we acquired confocal slices over 0.4 µm z-intervals. Due to technical limitations with SIM super-resolution imaging, we sampled the area of fluorescently labeled lysosomes by acquiring 3 confocal slices in the mid-point of the cell, where we quantified the pixel area for each slice and reported an average per cell. We then used Volocity 6.3.0 image analysis software to quantify the average number of fluorescent voxels or pixels within each cell. Due to the variation in lysosomal size from experiment to experiment we normalized the average voxel or pixel count to the corresponding control group. For lysosomal tubulation, we scored cells as positive for lysosome tubules if they displayed more than four lysosomal tubules greater than 4 μm. For antigen presentation analysis, we acquired confocal slices over 0.3 µm z-intervals and used Volocity to determine the total fluorescence intensity of antigen-MHCII complexes for 50-100 cells per experiment. To control for background, we established a threshold fluorescence intensity measure using a no-antigen control group for during each experiment. Image manipulation was done with ImageJ or Adobe Photoshop (Adobe Systems, San Jose, CA), without altering the relative signals within images or how data may be interpreted. All figures were assembled using Adobe Illustrator (Adobe Systems).

### T cell activation assays

The I-A^b^ restricted Eα-specific 1H3.1 T cell hybridoma cell-line was used for activation assays to recognize pre-activated dendritic cells expressing I-A^b^::Eα^52–68^ complexes. T-cells were cultured in RPMI-1640 medium supplemented with 10% heat-inactivated fetal bovine serum (Wisent) and 55 µM β-mercaptoethanol at 37°C, with 5% CO_2_. For activation assays, pre-activated DCs were mildly fixed in 1% PFA for 15 minutes at room temperature. Following fixation, cells were washed with PBS three times and then quenched thrice in complete medium for 10 minutes each, at room temperature. After, 1H3.1 T-cells and fixed-DCs were co-cultured at 2:1 and incubated for 40 hours at 37°C with 5% CO_2_. Next, the tissue culture medium was collected, and T-cells were immediately isolated following centrifugation at 800xg for 5 minutes. The supernatant was immediately stored in −80°C for downstream IL-2 secretion analysis. To quantify T-cell activation, secreted IL-2 samples were diluted 1:10 and subsequently analyzed using Mouse IL-2 Quantikine ELISA Kit (R&D Systems Inc, Minneapolis, MN), as per manufacturer’s specifications.

### Puromycylation and Western blotting

For puromycylation assays, cells were treated with 10 μg/mL of puromycin (Sigma-Aldrich), or an equivalent water volume for the non-puromycin group, for the last 15 min of each treatment. For all western blot analysis, cells were lysed in Laemmli buffer supplemented with 1:100 protease inhibitor cocktail (Sigma-Aldrich) and PhosSTOP protease inhibitor (Roche, Mississauga, ON) following each treatment. We loaded ∼0.8-1×10^6^ cell-equivalent per lane and proteins were then separated in a 10% or 15% SDS-PAGE, for high and low molecular weight proteins, respectively. Proteins were transferred to a polyvinylidene difluoride (PVDF) membrane (EMD Millipore, Toronto, ON), and blocked in 5% skim milk or BSA, in Tris-buffered saline buffer with 0.1% Tween 20 (TBST). Membranes were then immunoblotted using the appropriate primary and secondary antibodies prepared in 5% skim milk or BSA in TBST at the indicated dilutions. The primary antibodies used were rabbit anti-cathepsin D, ATP6V1H, ATP6V1D (GeneTex Inc., Irvine, CA), S6 ribosomal protein, phospho^Ser240/244^-S6 ribosomal protein, p70 S6 kinase, phospho^Thr389^-p70 S6 kinase, 4E-BP1, phospho^Thr37/46^-4E-BP, β-actin, Ha-Tag and Tata-box binding protein (TBP; Cell Signaling Technologies, Danvers, MA), all at 1:1,000. We also used mouse anti-puromycin clone 12D10 (1:1000, EMD Millipore), rat anti-LAMP1 (1:200; Developmental Studies Hybridoma Bank, Iowa City, IO) and secondary HRP-linked antibodies raised in donkey (1:10,000, Bethyl). Proteins were detected using Clarity enhanced chemiluminescence (Bio-Rad Laboratories, Mississauga, ON) with a ChemiDoc XRS+ or ChemiDoc Touch imaging system (Bio-Rad). Protein quantification was performed using Image Lab software (Bio-Rad), where protein loading was normalized to levels of Tata box binding protein (TBP) or β-actin, and then normalized against the vehicle group. Uncut and unedited images of the Western blots shown in each figure can be found in S1 Raw Images.

### LC3 conversion autophagy assay

We measured the conversion of LC3-I to LC3-II using immunoblotting, to measure effects on autophagy induction, in response to the pharmacological inhibitors used in our study. Primary macrophages were treated with the respective inhibitors as previously described. As a positive control for autophagy induction, we treated cells with Concanamycin A for 2 hours and/or cultured cells in Earle’s balanced salt solution (EBSS) (Gibco) for 2 and 6 hours. Cells were lysed with Laemmli buffer (as described previously) and processed for SDS-PAGE. Proteins were separated on a 20% poly-acrylamide gel using standard SDS-PAGE. Western blotting was performed as previously described, using a primary rabbit anti-LC3 antibody (1:1000; Cell Signalling Technologies) to detect LC3-I and LC3-II abundance. For autophagy induction, immunoblots were quantified where a ratio of LC3-II to LC3-I was determined and normalized to loading control actin. Relative autophagy index was determined by comparing each treatment group to resting cells.

### Quantitative RT-PCR

For RT-qPCR analysis in BMDMs, total RNA was extracted using the GeneJET RNA purification kit (ThermoFisher). Following RNA isolation, equal quantities of mRNA were reverse transcribed with iScript Reverse Transcription Super Mix (Bio-Rad) following manufacturer’s guidelines. The subsequent cDNA was amplified for quantitative PCR using the TaqMan Fast Advanced Master Mix (ThermoFisher) with appropriate TaqMan assays. The CFX96 Touch Real-Time PCR Detection System (Bio-Rad) and CFX Manager Software (Bio-Rad) were used for amplification and analysis. The TaqMan gene expression assays (ThermoFisher) for the reference genes Abt1 (Mm00803824_m1), B2M (Mm00437762_m1) and for target genes Atp6v1h (Mm00505548_m1), Atp6v1d (Mm00445832_m1), Lamp1 (Mm00495262_m1), Mcoln1 (Mm00522550_m1), CtsD (Mm00515586_m1), Lamp3/CD63 ((Mm01966817_g1), Lamp2 (Mm00495267_m1) and IL-6 (Mm00446190_m1) were done in triplicate. Target gene expression was determined by relative quantification (ΔΔCt method) to Abt1 and the vehicle-treated control sample.

### Polysome profiling

Polysome profiling was performed as detailed in Gandin *et al*. [77]. RAW264.7 cells were seeded in a 15-cm Petri dish and treated for 2 h or 6 h with a vehicle (DMSO), 500 ng/mL LPS from *Salmonella enterica* serotype minnesota Re 595, 100 nM torin1 for 2 h only, or the combination of LPS (500 ng/mL) and torin1 (100 nM) whereby cells were pre-treated for 15 minutes with torin1 before stimulation with LPS. Cells were harvested at 80% confluency, washed twice with ice-cold PBS containing 100 µg/mL cycloheximide and then lysed in hypotonic lysis buffer (5 mM Tris HCl, pH 7.5, 2.5 mM MgCl_2_, 1.5 mM KCl, 100 µg/ml cycloheximide, 2 mM dithiothreitol (DTT), 0.5% Triton, and 0.5% sodium deoxycholate). Optical density values at 260 nm (OD_260_) were measured in each lysate and 15 OD_260_ were then loaded on 5–50% sucrose gradients generated using Gradient Master (Biocomp, Fredericton, New Brunswick). Ten percent of lysates were saved as input samples for total RNA extraction. Sucrose gradients were subjected to ultracentrifugation (SW41 Ti 11E1698 rotor; Beckman at 260,000xg for 2 h at 4□°C) and fractionated by displacement using 60% sucrose/0.01% bromophenol blue on an ISCO Foxy fraction collector (35 s for each fraction, or ∼ 750□μL per fraction) equipped with an ultraviolet lamp for continuous absorbance monitoring at 254 nm. Fractions were flash-frozen immediately after fractionation and stored at −80□°C. RNA was isolated with Trizol (Thermofisher) as per manufacturer’s instruction. All experiments were carried out at least three independent biological replicates (n=3).

Reverse transcription and RT-qPCR were performed with iScript Reverse Transcription Super Mix (Bio-Rad) and TaqMan Fast Advanced Master Mix (ThermoFisher), respectively. All experiments were carried out at least three independent biological replicates (n=3). Analyses were carried out using relative standard curve method as instructed by the manufacturer. The following TaqMan assays were done using the primers described above for quantitative RT-PCR and in addition to NPC2 (Mm00499230_m1), Cln3 (Mm00487021_m1), Slc11a1 (Mm00443045_m1), Slc11a2 (Mm00435363_m1), CtsC (Mm00515580_m1), Sgsh (Mm00450747_m1), M6PR (Mm04208409_gH), Hyal1 (Mm01230688_g1), Actb (Mm02619580_g1) and Ppia (Mm02342430_g1).

### Global polysome profiling and analysis

RNA sequencing libraries were prepared using the Illumina TruSeq Stranded total RNA protocol including ribozero treatment (by the National Genomics Infrastructure, ScilifeLab, Stockholm, Sweden). Paired end sequencing was performed using NovaSeq6000 with control software 1.6.0/RTA v3.4.4. The resulting RNAseq reads were processed using the nextflow RNAseq pipeline (version 1.3; https://nf-co.re/) using default settings. Within the nextflow pipeline, high quality of sequencing reads was assured using fastQC (http://www.bioinformatics.babraham.ac.uk/projects/fastqc). Sequencing reads were then aligned to the GRCm38 genome using Hisat2 [109] followed by read summarization to assess expression levels using the featureCounts function [110] with Ensembl annotation [111]. Only protein coding genes localized to chromosomes 1 to 22, X, Y and MT were included. Genes with 0 counts in at least one sample were discarded. Raw counts were then analyzed using the anota2seq algorithm (version 1.4.2; [81]) with TMM-log2 normalization [112]. Analysis of changes in translation efficiencies, buffering, total mRNA and polysome-associated mRNA were performed using the *anota2seqAnalyze()* function. Changes were considered significant when passing the following parameters within the *anota2seqSelSigGenes()* function: maxPAdj = 0.25, minSlopeTranslation = −1, maxSlopeTranslation = 2, minSlopeBuffering = −2, maxSlopeBuffering = 1, selDeltaPT = log2(1.2), selDetaTP = log2(1.2), selDeltaP = 0 and selDeltaT = 0. Modes for regulation of gene expression were then determined using the *anota2seqRegModes()* function. The KEGG pathway database was used to extract genes annotated to the lysosome pathway [113, 114]. The RNAseq data is deposited on the Gene Expression Omnibus (GEO) with accession number GSE136470.

## Acknowledgments

We would like to thank Dr. Rosa Puertollano at the NIH for her kind donation of the CRISPR-deleted RAW strains (*tfeb*^−/−^, *tfe3*^−/−^ and *tfeb*^−/−^ *tfe3*^−/^ cells). We would like to thank Paul Paroutis and Michael Woodside from the Hospital for Sick Children Imaging Facility (Toronto, ON), and Christopher Spring from St. Michael’s Hospital Flow Cytometry Facility (Toronto, ON), for their technical advice and expertise. We would like to thank the technical support staff at the Vivarium Facilities at St. Michael’s Hospital for assistance in training and mice maintenance. The authors acknowledge assistance from the National Genomics Infrastructure in Genomics Production Stockholm supported by Science for Life Laboratory, the Knut and Alice Wallenberg Foundation and the Swedish Research Council, and SNIC/Uppsala Multidisciplinary Center for Advanced Computational Science for assistance with massively parallel sequencing and access to the UPPMAX computational infrastructure. The LAMP1 hybridoma antibody, developed by J.T. August, was obtained from the Developmental Studies Hybridoma Bank, created by the NICHD of the NIH and maintained at The University of Iowa, Department of Biology, Iowa City, IA.

## Supporting information

**S1 Fig: Preservation of tubules during fixation and super-resolution imaging.** (a) RAW macrophage lysosomes labeled with fluid phase fluorescent probes were imaged live or fixed with 4% PFA or a mixture of PFA and glutaraldehyde as explained in methods. (b) Percent lysosome tubulation was recorded within the population for cells exhibiting 4 or more lysosomal tubules longer than 4 µm. Statistical analysis was done with an ANOVA, where * indicates conditions that are statistically distinct from the corresponding resting group (*p<0.05). (c) Wide-field (WF) illumination or structured illumination microscopy (SIM) images of lysosomes in RAW macrophages, bone marrow derived macrophages (BMDM) and bone-marrow derived dendritic cells (BMDCs), before and after 2 h of LPS stimulation. Scale bar = 5 µm. See S9 Data for original data in S1 Fig.

**S2 Fig: Activated RAW macrophages have a larger lysosome holding capacity.** (a) Accumulation of Lucifer yellow (LY) in resting and activated RAW macrophages. RAW cells were stimulated and then allowed to internalize LY over time. (b) Pinocytosis rate by quantifying uptake of Lucifer yellow in RAW macrophages treated as indicated. (c) Retention of Lucifer yellow chased in probe-free medium in RAW cells previously treated as indicated and pre-labelled with Lucifer yellow for 1 h. In all cases, fluorescence measurements were done by flow cytometry. (d) Pinocytosis in increasingly maturing DCs exposed to LPS. Microscopy was used to measure the uptake of fluorescent dextran for 30 min by DCs exposed to LPS over indicated time points. Shown is the mean ± standard error of the mean from at least three experiments. For statistical analysis, ANOVA or Analysis of Covariance was used, whereby an asterisk indicates a significant difference in fluorescent probe levels compared to resting (*p<0.05). See S10 Data for original data in S2 Fig.

**S3 Fig: LPS increases lysosomal protein synthesis through mTOR and S6K.** (a) Western blot analysis of additional lysosomal proteins from whole cell lysates of resting primary macrophages or macrophages exposed to the indicated combinations and time of LPS, cycloheximide (CHX), Torin1, LY2584702 (LY2), AKT inhibitor (AKTi). (b) Quantification of Western blots showing the levels of LAMP2, TRPML1 and CD63 (LAMP3) normalized to actin. Data shown as the mean ± SEM from at least 3 independent experiments. For A and B, “2/4” indicates cells stimulated with 2 h of LPS, followed by a 4 h chase, whereas 2 and 6 h represent cells continuously exposed to LPS. See S11 Data for original data in S3 Fig.

**S4 Fig: Basal lysosome properties and trafficking is indistinguishable in wild-type RAWs and strains deleted for TFEB and/or TFE3.** (a-b) Western blot analysis of whole cell lysates from TFEB^−/−^, TFE3^−/−^ and double deleted cell-lines. (b) Quantification showing mutant lines are devoid of TFEB and/or TFE3 proteins, from three independent blots. (c) LAMP1 levels in whole cell lysates from wild-type and deletion mutants of TFEB and/or TFE3. (d) Quantification of LAMP1 levels in knock-out cells. LAMP1 levels were normalized to β-actin to control for loading. Statistical analysis using ANOVA determined that LAMP1 levels did not vary across strains. (e) Co-localization of dextran and LAMP1 in wild-type and deletion strains. Right, middle and left panels show dextran (red), endogenous LAMP1 (green) and merge, respectively. Scale bar = 5 µm. (f) Mander’s coefficient of dextran co-localizing in LAMP1 structures. Data shown as relative units (R.U), normalized to wild-type strain. (g) Pinocytosis label after a 1 hr pulse and 1 hr chase of fluorescent dextran in resting wild-type and deletion RAW strains, measured by microscopy and image analysis. Mean fluorescence intensity was normalized to wild-type strain and is represented as relative units (R.U). (h) Dextran fluorescence in RAW and deletion strains 2 h after LPS exposure or vehicle. For all data, shown are the mean± standard deviation from at least three independent experiments. See S12 Data for original data in S4 Fig.

**S5 Fig: LPS stimulates global protein synthesis through mTOR-S6K-4E-BP axes.** (a) Western blot analysis of whole cell lysates from resting and activated primary macrophages. Total levels and phosphorylation status of S6K and 4E-BP1 were monitored using the indicated antibodies. TBP served as a loading control. (b-c) Normalized ratio of (b) p-p70S6K and (c) p-4EBP1 to total p70S6K and 4E-BP1 protein. Shown is the mean ± standard deviation from three independent blots. (d) Western blot analysis of LC3-I to LC3-II conversion to measure treatment effect on autophagy induction in primary macrophages. BMDMs were activated with LPS in the presence or absence of protein synthesis, mTOR and S6K inhibitors for the time points indicated in brackets. Concanamycin A (ConA) and EBSS treatment was used as a positive control for autophagy induction. (e) Quantification of d from three independent experiments. Ratio of LC3II to LC3I levels was normalized to actin loading control. (f) Western blot analysis of protein puromycylation in resting and activated primary macrophages. LPS increases the amount of puromycylation indicating a boost in global protein synthesis that is blocked by mTOR inhibitors or cycloheximide. Lane 1 are control lysates from cells not exposed to puromycin. The band indicated by arrow is a non-specific band recognized by the anti-puromycin antibody. p-p70S6K and β-actin were used to monitor mTOR status and as a loading control, respectively. (g) Normalized puromycylation signal (excluding non-specific band) normalized over β-actin signal. Data is shown as the mean ± standard deviation from four independent experiments. For b, c, e, g statistical analysis was done with an ANOVA, where * or ** indicates conditions that are statistically distinct from control group (*p<0.05). (h) Normalized ratio of phosphorylated ribosomal S6 to total ribosomal S6 as depicted in Fig. 5f, in primary macrophages treated with LY2584702 (LY2) alone, or co-incubated with LPS for 2 h. Shown is the mean ± standard deviation of the mean from five independent blots. (i) Relative mRNA levels of select lysosomal genes (right) or interleukin-6 (left) in LPS and/or LY2 treated primary macrophages relative to Abt1 housekeeping gene and normalized against resting cells. Quantification was done with qRT-PCR by measuring the ÄÄCt as described in methods. Shown is the mean ± standard error of the mean from four independent experiments. See S13 Data for original data in S5 Fig.

**S6 Fig: Polysome profiling of RAW macrophages: additional replicate data.** Percent of target mRNA (a: LAMP1, b: ATP6V1H, c: ATP6V1D, d: CtsD, e: β-actin, f: PPIA, and g: B2M) associated with each ribosome fraction in resting, LPS-treated macrophages and macrophages co-exposed to LPS and torin1, or treated with torin1 alone. Left, middle and right panels show 2 h, 6 h and torin1 (2 h) treatments, respectively. Shown, is an additional biological replicate of the experiment described in Fig 7. See S14 Data for original data in S6 Fig.

**S7 Fig: The effects of RAW macrophage stimulation by LPS on protein synthesis**. Percent of target mRNA (a: LAMP1, b: ATP6V1H, c: ATP6V1D, d: CtsD, e: β-actin, f: PPIA, and g: B2M) associated with each ribosome fraction in resting and torin1 (2 h; 100 nM) treated cells for data presented in Fig 7. See S15 Data for original data in S7 Fig.

**S8 Fig: Polysome profiling validation of select targets identified through RNA-Seq analysis.** Percent of target mRNA (a: NPC2, b: Cln3, c: Slc11a1, d: CtsC, e: Slc11a2, f: Sgsh, g: M6PR, and h: Hyal1) associated with each ribosome fraction in resting, LPS-treated macrophages and macrophages co-exposed to LPS and torin1. See S16 Data for original data in S8 Fig.

**S9 Fig: Polysome profiling of two biological replicates used for global RNAseq analysis in Fig 8.** Percent of target mRNA (a: LAMP1, b: ATP6V1H, c: ATP6V1D, d: β-actin, e: PPIA, and f: B2M) associated with each polysome fraction in resting, LPS-treated macrophages and macrophages co-exposed to LPS and torin1 for 6 hours. Biological replicate 1 (left) and replicate 2 (right) of data presented in Fig 8 global RNAseq analysis from a total of three experiments. See S17 Data for original data in S9 Fig.

**S10 Fig: Effect of LY2584702 and Torin1 treatments on HEL presentation by BMDCs.** (a) I-A^k^::HEL^46-61^ presentation in BMDCs after incubation with HEL for 6 hours in the presence and/or absence of LPS, torin1 and LY2. I-A^k^::HEL^46-61^ cell surface levels were detected by staining unpermeabilized cells with the monoclonal antibody Aw3.18.14. (b) Quantification of total average fluorescence intensity of I-A^k^::HEL^46-61^ complexes at the plasma membrane. Shown is the mean ± SD from three experiments, where 50-100 cells were quantified for each. Data was analyzed using ANOVA, whereby * indicates a difference compared to the Resting + HEL condition and ** indicates a difference compared to HEL+LPS (p<0.05). Scale bar = 15µm. Colour scale: 0 – 12000 (low-high). See S18 Data for original data in S10 Fig.

**S11 Fig: A model for mTORC1-dependent regulation of lysosome remodeling in phagocytes in response to LPS stimulation.** LPS engages the PI3K-AKT-mTOR signal axis to stimulate mTORC1 activity. We suggest that mTORC1 then regulates two parallel pathways to modulate lysosome size and morphology: i) mTORC1 activity augments Arl8b GTPase levels on the lysosome membrane to boost kinesin-1 recruitment to coordinate lysosome extension and anterograde transport. ii) in parallel, mTORC1 stimulates S6Ks and inhibits 4E-BPs to promote translation and rapidly boost levels of various (select) endo-lysosomal proteins, catalyzing endo-lysosome expansion. This expansion increases the holding capacity of the endo-lysosomal system, likely promoting antigen retention. Together both pathways (i and ii) converge to promote lysosome remodeling, collectively bolstering immunity. Importantly, this model does not imply that mTORC1 has no additional functions contributing to phagocyte activation and antigen presentation, nor does it imply that enhanced translation only boosts endo-lysosomal function.

**S1 Table: Original data for the RNAseq global analysis in Fig 8.**

**S1 Data: Original data represented in Fig 1.**

**S2 Data: Original data represented in Fig 2.**

**S3 Data: Original data represented in Fig 3.**

**S4 Data: Original data represented in Fig 4.**

**S5 Data: Original data represented in Fig 5.**

**S6 Data: Original data represented in Fig 6.**

**S7 Data: Original data represented in Fig 7.**

**S8 Data: Original data represented in Fig 9.**

**S9 Data: Original data represented in S1 Fig.**

**S10 Data: Original data represented in S2 Fig.**

**S11 Data: Original data represented in S3 Fig.**

**S12 Data: Original data represented in S4 Fig.**

**S13 Data: Original data represented in S5 Fig.**

**S14 Data: Original data represented in S6 Fig.**

**S15 Data: Original data represented in S7 Fig.**

**S16 Data: Original data represented in S8 Fig.**

**S17 Data: Original data represented in S9 Fig.**

**S18 Data: Original data represented in S10 Fig.**

**S1_Raw_images: Unedited, original western blots images**.av

## Notes

#### Summary of Updates

New data to address peer review done including Figure 8 (RNA-seq). T cell tests were added.

## References

1. Saffi GT, Botelho RJ. Lysosome Fission: Planning for an Exit [Internet]. Trends in Cell Biology. 2019. pp. 635–646. doi:10.1016/j.tcb.2019.05.003

2. Bright NA, Davis LJ, Luzio JP. Endolysosomes Are the Principal Intracellular Sites of Acid Hydrolase Activity. Curr Biol. Elsevier; 2016;26: 2233–45. doi:10.1016/j.cub.2016.06.046

3. Chan Y-HM, Reyes L, Sohail SM, Tran NK, Marshall WF. Organelle Size Scaling of the Budding Yeast Vacuole by Relative Growth and Inheritance. Curr Biol. 2016;26: 1221–1228. doi:10.1016/j.cub.2016.03.020

4. Mullins C, Bonifacino JS. The molecular machinery for lysosome biogenesis [Internet]. BioEssays. John Wiley & Sons, Inc.; 2001. pp. 333–343. doi:10.1002/bies.1048

5. Behnia R, Munro S. Organelle identity and the signposts for membrane traffic. Nature. 2005;438: 597–604. doi:10.1038/nature04397

6. Levy DL, Heald R. Mechanisms of Intracellular Scaling. Annu Rev Cell Dev Biol. 2012;28: 113–135. doi:10.1146/annurev-cellbio-092910-154158

7. Mills JC, Taghert PH. Scaling factors: Transcription factors regulating subcellular domains. BioEssays. 2012;34: 10–16. doi:10.1002/bies.201100089

8. Porta C, Riboldi E, Ippolito A, Sica A. Molecular and epigenetic basis of macrophage polarized activation [Internet]. Seminars in Immunology. 2015. pp. 237–248. doi:10.1016/j.smim.2015.10.003

9. Trombetta ES, Ebersold M, Garrett W, Pypaert M, Mellman I. Activation of lysosomal function during dendritic cell maturation. Science (80-). 2003;299: 1400–1403. doi:10.1126/science.1080106

10. Kelly B, O’Neill LAJ. Metabolic reprogramming in macrophages and dendritic cells in innate immunity. Cell Res. 2015;25: 771–84. doi:10.1038/cr.2015.68

11. Janssens S, Pulendran B, Lambrecht BN. Emerging functions of the unfolded protein response in immunity. Nat Immunol. 2014;15: 910–919. doi:10.1038/ni.2991

12. Hipolito VEB, Ospina-Escobar E, Botelho RJ. Lysosome remodelling and adaptation during phagocyte activation [Internet]. Cellular Microbiology. 2018. p. e12824. doi:10.1111/cmi.12824

13. Delamarre L, Pack M, Chang H, Mellman I, Trombetta ES. Differential lysosomal proteolysis in antigen-presenting cells determines antigen fate. Science. 2005;307: 1630–4. doi:10.1126/science.1108003

14. Gray MA, Choy CH, Dayam RM, Ospina-Escobar E, Somerville A, Xiao X, et al. Phagocytosis Enhances Lysosomal and Bactericidal Properties by Activating the Transcription Factor TFEB. Curr Biol. 2016;26. doi:10.1016/j.cub.2016.05.070

15. Mrakovic A, Kay JG, Furuya W, Brumell JH, Botelho RJ. Rab7 and Arl8 GTPases are Necessary for Lysosome Tubulation in Macrophages. Traffic. 2012;13: 1667–1679. doi:10.1111/tra.12003

16. Vyas JM, Kim Y-M, Artavanis-Tsakonas K, Love JC, Van der Veen AG, Ploegh HL. Tubulation of class II MHC compartments is microtubule dependent and involves multiple endolysosomal membrane proteins in primary dendritic cells. J Immunol. 2007;178: 7199–210.

17. Saric A, Hipolito VEB, Kay JG, Canton J, Antonescu CN, Botelho RJ. mTOR controls lysosome tubulation and antigen presentation in macrophages and dendritic cells. Mol Biol Cell. 2016;27: 321–333. doi:10.1091/mbc.E15-05-0272

18. Hollenbeck PJ, Swanson JA. Radial extension of macrophage tubular lysosomes supported by kinesin. Nature. 1990;346: 864–866. doi:10.1038/346864a0

19. Li X, Rydzewski N, Hider A, Zhang X, Yang J, Wang W, et al. A molecular mechanism to regulate lysosome motility for lysosome positioning and tubulation. Nat Cell Biol. 2016;18: 404–17. doi:10.1038/ncb3324

20. Swanson J, Burke E, Silverstein SC. Tubular lysosomes accompany stimulated pinocytosis in macrophages. J Cell Biol. 1987;104: 1217–1222. doi:10.1083/jcb.104.5.1217

21. Chow A, Toomre D, Garrett W, Mellman I. Dendritic cell maturation triggers retrograde MHC class II transport from lysosomes to the plasma membrane. Nature. 2002;418: 988– 94. doi:10.1038/nature01006

22. Boes M, Cerny J, Massol R, Op den Brouw M, Kirchhausen T, Chen J, et al. T-cell engagement of dendritic cells rapidly rearranges MHC class II transport. Nature. 2002;418: 983–8. doi:10.1038/nature01004

23. Nakamura N, Lill JR, Phung Q, Jiang Z, Bakalarski C, de Mazière A, et al. Endosomes are specialized platforms for bacterial sensing and NOD2 signalling. Nature. 2014;509: 240– 4. doi:10.1038/nature13133

24. Mantegazza AR, Zajac AL, Twelvetrees A, Holzbaur ELF, Amigorena S, Marks MS. TLR-dependent phagosome tubulation in dendritic cells promotes phagosome cross-talk to optimize MHC-II antigen presentation. Proc Natl Acad Sci U S A. 2014;111: 15508–13. doi:10.1073/pnas.1412998111

25. Mony VK, Benjamin S, O’Rourke EJ. A lysosome-centered view of nutrient homeostasis. Autophagy. 2016;12: 619–631. doi:10.1080/15548627.2016.1147671

26. Jewell JL, Russell RC, Guan K-L. Amino acid signalling upstream of mTOR. Nat Rev Mol Cell Biol. 2013;14: 133–9. doi:10.1038/nrm3522

27. Lim CY, Zoncu R. The lysosome as a command-and-control center for cellular metabolism. J Cell Biol. 2016;214: 653–664. doi:10.1083/jcb.201607005

28. Inpanathan S, Botelho RJ. The Lysosome Signaling Platform: Adapting With the Times. Front Cell Dev Biol. 2019;7: 113. doi:10.3389/fcell.2019.00113

29. Sancak Y, Bar-Peled L, Zoncu R, Markhard AL, Nada S, Sabatini DM. Ragulator-rag complex targets mTORC1 to the lysosomal surface and is necessary for its activation by amino acids. Cell. 2010;141: 290–303. doi:10.1016/j.cell.2010.02.024

30. Zoncu R, Bar-Peled L, Efeyan A, Wang S, Sancak Y, Sabatini DM. mTORC1 senses lysosomal amino acids through an inside-out mechanism that requires the vacuolar H+-ATPase. Science (80-). 2011;334: 678–683. doi:10.1126/science.1207056

31. Efeyan A, Zoncu R, Sabatini DM. Amino acids and mTORC1: From lysosomes to disease [Internet]. Trends in Molecular Medicine. 2012. pp. 524–533. doi:10.1016/j.molmed.2012.05.007

32. Martina JA, Puertollano R. Rag GTPases mediate amino acid-dependent recruitment of TFEB and MITF to lysosomes. J Cell Biol. 2013;200: 475–491. doi:10.1083/jcb.201209135

33. Bar-Peled L, Schweitzer LD, Zoncu R, Sabatini DM. Ragulator is a GEF for the rag GTPases that signal amino acid levels to mTORC1. Cell. Elsevier Inc.; 2012;150: 1196– 1208. doi:10.1016/j.cell.2012.07.032

34. Zhang CS, Jiang B, Li M, Zhu M, Peng Y, Zhang YL, et al. The lysosomal v-ATPase-ragulator complex is a common activator for AMPK and mTORC1, acting as a switch between catabolism and anabolism. Cell Metab. 2014;20: 526–540. doi:10.1016/j.cmet.2014.06.014

35. Zoncu R, Efeyan A, Sabatini DM. MTOR: From growth signal integration to cancer, diabetes and ageing. Nature Reviews Molecular Cell Biology. 2011. pp. 21–35. doi:10.1038/nrm3025

36. Thoreen CC. The molecular basis of mTORC1-regulated translation. Biochem Soc Trans. 2017;45: 213–221. doi:10.1042/BST20160072

37. Buszczak M, Signer RAJ, Morrison SJ. Cellular differences in protein synthesis regulate tissue homeostasis [Internet]. Cell. 2014. pp. 242–251. doi:10.1016/j.cell.2014.09.016

38. Magnuson B, Ekim B, Fingar DC. Regulation and function of ribosomal protein S6 kinase (S6K) within mTOR signalling networks. Biochem J. 2012;441: 1–21. doi:10.1042/BJ20110892

39. Roux PP, Topisirovic I. Signaling pathways involved in the regulation of mRNA translation. Mol Cell Biol. 2018;38: MCB.00070-18. doi:10.1128/MCB.00070-18

40. Holz MK, Ballif BA, Gygi SP, Blenis J. mTOR and S6K1 mediate assembly of the translation preinitiation complex through dynamic protein interchange and ordered phosphorylation events. Cell. 2005;123: 569–580. doi:10.1016/j.cell.2005.10.024

41. Burnett PE, Barrow RK, Cohen NA, Snyder SH, Sabatini DM. RAFT1 phosphorylation of the translational regulators p70 S6 kinase and 4E-BP1. Proc Natl Acad Sci U S A. 1998;95: 1432–7.

42. Beretta L, Gingras AC, Svitkin Y V, Hall MN, Sonenberg N. Rapamycin blocks the phosphorylation of 4E-BP1 and inhibits cap-dependent initiation of translation. EMBO J. 1996;15: 658–64.

43. Jung CH, Jun CB, Ro S-H, Kim Y-M, Otto NM, Cao J, et al. ULK-Atg13-FIP200 Complexes Mediate mTOR Signaling to the Autophagy Machinery. Mol Biol Cell. 2009;20: 1992–2003. doi:10.1091/mbc.E08-12-1249

44. Ganley IG, Lam DH, Wang J, Ding X, Chen S, Jiang X. ULK1·ATG13·FIP200 complex mediates mTOR signaling and is essential for autophagy. J Biol Chem. 2009;284: 12297– 12305. doi:10.1074/jbc.M900573200

45. Settembre C, Zoncu R, Medina DL, Vetrini F, Erdin SS, Erdin SS, et al. A lysosome-to-nucleus signalling mechanism senses and regulates the lysosome via mTOR and TFEB. Eur Mol Biol Organ J. Nature Publishing Group; 2012;31: 1095–108. doi:10.1038/emboj.2012.32

46. Roczniak-Ferguson A, Petit CS, Froehlich F, Qian S, Ky J, Angarola B, et al. The transcription factor TFEB links mTORC1 signaling to transcriptional control of lysosome homeostasis. Sci Signal. 2012;5: ra42. doi:10.1126/scisignal.2002790

47. Owen KA, Meyer CB, Bouton AH, Casanova JE. Activation of Focal Adhesion Kinase by Salmonella Suppresses Autophagy via an Akt/mTOR Signaling Pathway and Promotes Bacterial Survival in Macrophages. Deretic V, editor. PLoS Pathog. Public Library of Science; 2014;10: e1004159. doi:10.1371/journal.ppat.1004159

48. Moon J-S, Hisata S, Park M-A, DeNicola GM, Ryter SW, Nakahira K, et al. mTORC1-Induced HK1-Dependent Glycolysis Regulates NLRP3 Inflammasome Activation. Cell Rep. 2015;12: 102–115. doi:10.1016/j.celrep.2015.05.046

49. Lelouard H, Schmidt EK, Camosseto V, Clavarino G, Ceppi M, Hsu HT, et al. Regulation of translation is required for dendritic cell function and survival during activation. J Cell Biol. The Rockefeller University Press; 2007;179: 1427–1439. doi:10.1083/jcb.200707166

50. Pan H, Zhong XP, Lee S. Sustained activation of mTORC1 in macrophages increases AMPKα-dependent autophagy to maintain cellular homeostasis. BMC Biochem. 2016;17: 14. doi:10.1186/s12858-016-0069-6

51. Terawaki S, Camosseto V, Prete F, Wenger T, Papadopoulos A, Rondeau C, et al. RUN and FYVE domain-containing protein 4 enhances autophagy and lysosome tethering in response to Interleukin-4. J Cell Biol. Rockefeller University Press; 2015;210: 1133– 1152. doi:10.1083/jcb.201501059

52. Pan H, O’Brien TF, Wright G, Yang J, Shin J, Wright KL, et al. Critical Role of the Tumor Suppressor Tuberous Sclerosis Complex 1 in Dendritic Cell Activation of CD4 T Cells by Promoting MHC Class II Expression via IRF4 and CIITA. J Immunol. 2013;191: 699–707. doi:10.4049/jimmunol.1201443

53. Long F, Zhou J, Peng H. Visualization and analysis of 3D microscopic images. Lewitter F, editor. PLoS Comput Biol. Public Library of Science; 2012;8: e1002519. doi:10.1371/journal.pcbi.1002519

54. Walter T, Shattuck DW, Baldock R, Bastin ME, Carpenter AE, Duce S, et al. Visualization of image data from cells to organisms [Internet]. Nature Methods. NIH Public Access; 2010. pp. S26–S41. doi:10.1038/nmeth.1431

55. Gustafsson MGL. Nonlinear structured-illumination microscopy: Wide-field fluorescence imaging with theoretically unlimited resolution. Proc Natl Acad Sci. National Academy of Sciences; 2005;102: 13081–13086. doi:10.1073/pnas.0406877102

56. Barois N, de Saint-Vis B, Lebecque S, Geuze HJ, Kleijmeer MJ. MHC class II compartments in human dendritic cells undergo profound structural changes upon activation. Traffic. 2002;3: 894–905. doi:10.1034/j.1600-0854.2002.31205.x

57. Garrett WS, Chen LM, Kroschewski R, Ebersold M, Turley S, Trombetta S, et al. Developmental control of endocytosis in dendritic cells by Cdc42. Cell. 2000;102: 325–34.

58. Platt CD, Ma JK, Chalouni C, Ebersold M, Bou-Reslan H, Carano RAD, et al. Mature dendritic cells use endocytic receptors to capture and present antigens. Proc Natl Acad Sci. National Academy of Sciences; 2010;107: 4287–4292. doi:10.1073/pnas.0910609107

59. Drutman SB, Trombetta ES. Dendritic cells continue to capture and present antigens after maturation in vivo. J Immunol. 2010;185: 2140–6. doi:10.4049/jimmunol.1000642

60. Kobayashi T, Tanaka T, Toyama-Sorimachi N. How do cells optimize luminal environments of endosomes/lysosomes for efficient inflammatory responses. J Biochem. 2013;154: 491–499. doi:10.1093/jb/mvt099

61. Swanson J a, Yirinec BD, Silverstein SC. Phorbol esters and horseradish peroxidase stimulate pinocytosis and redirect the flow of pinocytosed fluid in macrophages. J Cell Biol. 1985;100: 851–859. doi:10.1083/jcb.100.3.851

62. Sardiello M, Palmieri M, di Ronza A, Medina DL, Valenza M, Gennarino VA, et al. A Gene Network Regulating Lysosomal Biogenesis and Function. Science (80-). 2009;325: 473–7. doi:10.1126/science.1174447

63. Settembre C, Fraldi A, Medina DL, Ballabio A. Signals from the lysosome: a control centre for cellular clearance and energy metabolism. Nat Rev Mol Cell Biol. Nature Publishing Group; 2013;14: 283–296. doi:10.1038/nrm3565

64. Martina JA, Diab HI, Lishu L, Jeong-A L, Patange S, Raben N, et al. The nutrient-responsive transcription factor TFE3 promotes autophagy, lysosomal biogenesis, and clearance of cellular debris. Sci Signal. 2014;7: ra9. doi:10.1126/scisignal.2004754

65. Polito V a, Li H, Martini-Stoica H, Wang B, Yang L, Xu Y, et al. Selective clearance of aberrant tau proteins and rescue of neurotoxicity by transcription factor EB. EMBO Mol Med. 2014;6: 1–19. doi:10.15252/emmm.201303671

66. Pastore N, Brady OA, Diab HI, Martina JA, Sun L, Huynh T, et al. TFEB and TFE3 cooperate in the regulation of the innate immune response in activated macrophages. Autophagy. 2016;12: 1240–1258. doi:10.1080/15548627.2016.1179405

67. Raben N, Puertollano R. TFEB and TFE3: Linking Lysosomes to Cellular Adaptation to Stress. Annu Rev Cell Dev Biol. 2016;32: 255–278. doi:10.1146/annurev-cellbio-111315-125407

68. Liu AP, Botelho RJ, Antonescu CN. The big and intricate dreams of little organelles: Embracing complexity in the study of membrane traffic. Traffic. 2017; doi:10.1111/tra.12497

69. Zhang X, Cheng X, Yu L, Yang J, Calvo R, Patnaik S, et al. MCOLN1 is a ROS sensor in lysosomes that regulates autophagy. Nat Commun. 2016;7: 12109. doi:10.1038/ncomms12109

70. Schott J, Reitter S, Philipp J, Haneke K, Schäfer H, Stoecklin G. Translational Regulation of Specific mRNAs Controls Feedback Inhibition and Survival during Macrophage Activation. Wells CA, editor. PLoS Genet. Public Library of Science; 2014;10: e1004368. doi:10.1371/journal.pgen.1004368

71. Graczyk D, White RJ, Ryan KM. Involvement of RNA Polymerase III in Immune Responses. Mol Cell Biol. American Society for Microbiology; 2015;35: 1848–1859. doi:10.1128/MCB.00990-14

72. Ivanov SS, Roy CR. Pathogen signatures activate a ubiquitination pathway that modulates the function of the metabolic checkpoint kinase mTOR. Nat Immunol. NIH Public Access; 2013;14: 1219–1228. doi:10.1038/ni.2740

73. Weichhart T, Costantino G, Poglitsch M, Rosner M, Zeyda M, Stuhlmeier KM, et al. The TSC-mTOR Signaling Pathway Regulates the Innate Inflammatory Response. Immunity. 2008;29: 565–577. doi:10.1016/j.immuni.2008.08.012

74. Schmitz F, Heit A, Dreher S, Eisenächer K, Mages J, Haas T, et al. Mammalian target of rapamycin (mTOR) orchestrates the defense program of innate immune cells. Eur J Immunol. John Wiley & Sons, Ltd; 2008;38: 2981–2992. doi:10.1002/eji.200838761

75. Schmidt EK, Clavarino G, Ceppi M, Pierre P. SUnSET, a nonradioactive method to monitor protein synthesis. Nat Methods. 2009;6: 275–277. doi:10.1038/nmeth.1314

76. Rong L, Livingstone M, Sukarieh R, Petroulakis E, Gingras A-C, Crosby K, et al. Control of eIF4E cellular localization by eIF4E-binding proteins, 4E-BPs. RNA. 2008;14: 1318–27. doi:10.1261/rna.950608

77. Gandin V, Sikström K, Alain T, Morita M, McLaughlan S, Larsson O, et al. Polysome fractionation and analysis of mammalian translatomes on a genome-wide scale. J Vis Exp. 2014; doi:10.3791/51455

78. Martina JA, Chen Y, Gucek M, Puertollano R. MTORC1 functions as a transcriptional regulator of autophagy by preventing nuclear transport of TFEB. Autophagy. 2012;8: 903–14. doi:10.4161/auto.19653

79. Piehler AP, Grimholt RM, Ovstebo R, Berg JP. Gene expression results in lipopolysaccharide-stimulated monocytes depend significantly on the choice of reference genes. BMC Immunol. BioMed Central; 2010;11: 21. doi:10.1186/1471-2172-11-21

80. Gordon EB, Hart GT, Tran TM, Waisberg M, Akkaya M, Skinner J, et al. Inhibiting the mammalian target of rapamycin blocks the development of experimental cerebral malaria. MBio. American Society for Microbiology; 2015;6: e00725. doi:10.1128/mBio.00725-15

81. Oertlin C, Lorent J, Murie C, Furic L, Topisirovic I, Larsson O. Generally applicable transcriptome-wide analysis of translation using anota2seq. Nucleic Acids Res. Narnia; 2019;47: e70–e70. doi:10.1093/nar/gkz223

82. Gandin V, Masvidal L, Hulea L, Gravel SP, Cargnello M, McLaughlan S, et al. NanoCAGE reveals 5’ UTR features that define specific modes of translation of functionally related MTOR-sensitive mRNAs. Genome Res. 2016;26: 636–648. doi:10.1101/gr.197566.115

83. Masvidal L, Hulea L, Furic L, Topisirovic I, Larsson O. mTOR-sensitive translation: Cleared fog reveals more trees. RNA Biol. 2017;14: 1299–1305. doi:10.1080/15476286.2017.1290041

84. Dadaglio G, Nelson CA, Deck MB, Petzold SJ, Unanue ER. Characterization and quantitation of peptide-MHC complexes produced from hen egg lysozyme using a monoclonal antibody. Immunity. 1997;6: 727–738. doi:10.1016/S1074-7613(00)80448-3

85. Murphy DB, Rath S, Pizzo E, Rudensky AY, George A, Larson JK, et al. Monoclonal antibody detection of a major self peptide. MHC class II complex. J Immunol. 1992;148: 3483–3491.

86. Viret C, Janeway CA. Functional and phenotypic evidence for presentation of E alpha 52-68 structurally related self-peptide(s) in I-E alpha-deficient mice. J Immunol. 2000;164: 4627–34.

87. Su X, Yu Y, Zhong Y, Giannopoulou EG, Hu X, Liu H, et al. Interferon-γ regulates cellular metabolism and mRNA translation to potentiate macrophage activation. Nat Immunol. 2015;16: 838–849. doi:10.1038/ni.3205

88. Piccirillo CA, Bjur E, Topisirovic I, Sonenberg N, Larsson O. Translational control of immune responses: From transcripts to translatomes [Internet]. Nature Immunology. 2014. pp. 503–511. doi:10.1038/ni.2891

89. Yates RM, Hermetter A, Taylor GA, Russell DG. Macrophage Activation Downregulates the Degradative Capacity of the Phagosome. Traffic. John Wiley & Sons, Ltd (10.1111); 2007;8: 241–250. doi:10.1111/j.1600-0854.2006.00528.x

90. Boes M, Bertho N, Cerny J, Op den Brouw M, Kirchhausen T, Ploegh H. T Cells Induce Extended Class II MHC Compartments in Dendritic Cells in a Toll-Like Receptor-Dependent Manner. J Immunol. American Association of Immunologists; 2003;171: 4081–4088. doi:10.4049/jimmunol.171.8.4081

91. Wessling-Resnick M. Nramp1 and other transporters involved in metal withholding during infection. J Biol Chem. 2015;290: 18984–18990. doi:10.1074/jbc.R115.643973

92. Settembre C, De Cegli R, Mansueto G, Saha PK, Vetrini F, Visvikis O, et al. TFEB controls cellular lipid metabolism through a starvation-induced autoregulatory loop. Nat Cell Biol. 2013;15: 647–658. doi:10.1038/ncb2718

93. Chua JP, Reddy SL, Merry DE, Adachi H, Katsuno M, Sobue G, et al. Transcriptional activation of TFEB/ZKSCAN3 target genes underlies enhanced autophagy in spinobulbar muscular atrophy. Hum Mol Genet. 2014;23: 1376–1386. doi:10.1093/hmg/ddt527

94. Thoreen CC, Chantranupong L, Keys HR, Wang T, Gray NS, Sabatini DM. A unifying model for mTORC1-mediated regulation of mRNA translation. Nature. 2012;485: 109– 113. doi:10.1038/nature11083

95. Nandagopal N, Roux PP. Regulation of global and specific mRNA translation by the mTOR signaling pathway. Transl (Austin, Tex). Taylor & Francis; 2015;3: e983402. doi:10.4161/21690731.2014.983402

96. Ma XM, Yoon S-O, Richardson CJ, Jülich K, Blenis J. SKAR Links Pre-mRNA Splicing to mTOR/S6K1-Mediated Enhanced Translation Efficiency of Spliced mRNAs. Cell. 2008;133: 303–313. doi:10.1016/j.cell.2008.02.031

97. Yuan S, Li J, Diener DR, Choma MA, Rosenbaum JL, Sun Z. Target-of-rapamycin complex 1 (Torc1) signaling modulates cilia size and function through protein synthesis regulation. Proc Natl Acad Sci U S A. National Academy of Sciences; 2012;109: 2021–6. doi:10.1073/pnas.1112834109

98. Meyuhas O. Synthesis of the translational apparatus is regulated at the translational level. Eur J Biochem. 2000;267: 6321–30.

99. Hsieh AC, Liu Y, Edlind MP, Ingolia NT, Janes MR, Sher A, et al. The translational landscape of mTOR signalling steers cancer initiation and metastasis. Nature. 2012;485: 55–61. doi:10.1038/nature10912

100. Masvidal L, Hulea L, Furic L, Topisirovic I, Larsson O. mTOR-sensitive translation: Cleared fog reveals more trees [Internet]. RNA Biology. 2017. pp. 1299–1305. doi:10.1080/15476286.2017.1290041

101. Yamashita R, Suzuki Y, Takeuchi N, Wakaguri H, Ueda T, Sugano S, et al. Comprehensive detection of human terminal oligo-pyrimidine (TOP) genes and analysis of their characteristics. Nucleic Acids Res. Oxford University Press; 2008;36: 3707–3715. doi:10.1093/nar/gkn248

102. Bilanges B, Argonza-Barrett R, Kolesnichenko M, Skinner C, Nair M, Chen M, et al. Tuberous sclerosis complex proteins 1 and 2 control serum-dependent translation in a TOP-dependent and -independent manner. Mol Cell Biol. 2007;27: 5746–5764. doi:10.1128/MCB.02136-06

103. Larsson O, Morita M, Topisirovic I, Alain T, Blouin M-J, Pollak M, et al. Distinct perturbation of the translatome by the antidiabetic drug metformin. Proc Natl Acad Sci. 2012;109: 8977–8982. doi:10.1073/pnas.1201689109

104. Rosa-Ferreira C, Munro S. Arl8 and SKIP Act Together to Link Lysosomes to Kinesin-1. Dev Cell. 2011;21: 1171–1178. doi:10.1016/j.devcel.2011.10.007

105. Michelet X, Garg S, Wolf BJ, Tuli A, Ricciardi-Castagnoli P, Brenner MB. MHC class II presentation is controlled by the lysosomal small GTPase, Arl8b. J Immunol. 2015;194: 2079–88. doi:10.4049/jimmunol.1401072

106. Liberman R, Bond S, Shainheit MG, Stadecker MJ, Forgac M. Regulated Assembly of Vacuolar ATPase Is Increased during Cluster Disruption-induced Maturation of Dendritic Cells through a Phosphatidylinositol 3-Kinase/mTOR-dependent Pathway. J Biol Chem. 2014;289: 1355–1363. doi:10.1074/jbc.M113.524561

107. Inaba K, Inaba M, Romani N, Aya H, Deguchi M, Ikehara S, et al. Generation of large numbers of dendritic cells from mouse bone marrow cultures supplemented with granulocyte/macrophage colony-stimulating factor. J Exp Med. 1992;176: 1693–1702. doi:10.1084/jem.176.6.1693

108. Weischenfeldt J, Porse B. Bone Marrow-Derived Macrophages (BMM): Isolation and Applications. Cold Spring Harb Protoc. 2008;2008: pdb.prot5080-pdb.prot5080. doi:10.1101/pdb.prot5080

109. Kim D, Langmead B, Salzberg SL. HISAT: A fast spliced aligner with low memory requirements. Nat Methods. Nature Publishing Group; 2015;12: 357–360. doi:10.1038/nmeth.3317

110. Liao Y, Smyth GK, Shi W. FeatureCounts: An efficient general purpose program for assigning sequence reads to genomic features. Bioinformatics. 2014;30: 923–930. doi:10.1093/bioinformatics/btt656

111. Kersey PJ, Allen JE, Allot A, Barba M, Boddu S, Bolt BJ, et al. Ensembl Genomes 2018: An integrated omics infrastructure for non-vertebrate species. Nucleic Acids Res. Narnia; 2018;46: D802–D808. doi:10.1093/nar/gkx1011

112. Robinson MD, McCarthy DJ, Smyth GK. edgeR: A Bioconductor package for differential expression analysis of digital gene expression data. Bioinformatics. Narnia; 2009;26: 139–140. doi:10.1093/bioinformatics/btp616

113. Kanehisa M, Sato Y, Furumichi M, Morishima K, Tanabe M. New approach for understanding genome variations in KEGG. Nucleic Acids Res. 2019;47: D590–D595. doi:10.1093/nar/gky962

114. Kanehisa M, Goto S. KEGG: Kyoto encyclopedia of genes and genomes. Nucleic Acids Res. 2000;28: 27–30. doi:10.1093/nar/28.1.27

