## Supplementary figures and images for "Enhanced translation expands the endo-lysosome size and promotes antigen presentation during phagocyte activation"

### S1 Fig

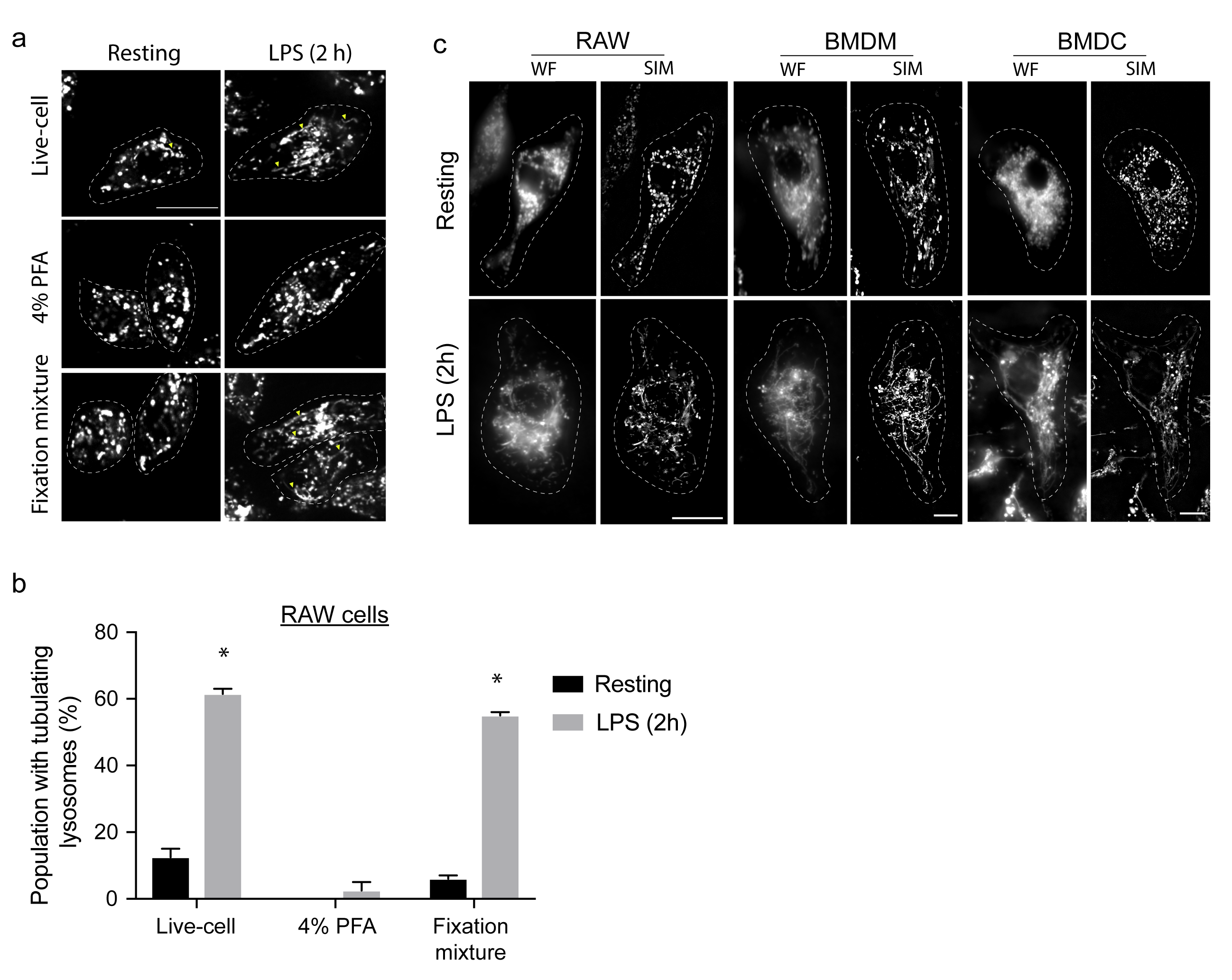

### S2 Fig

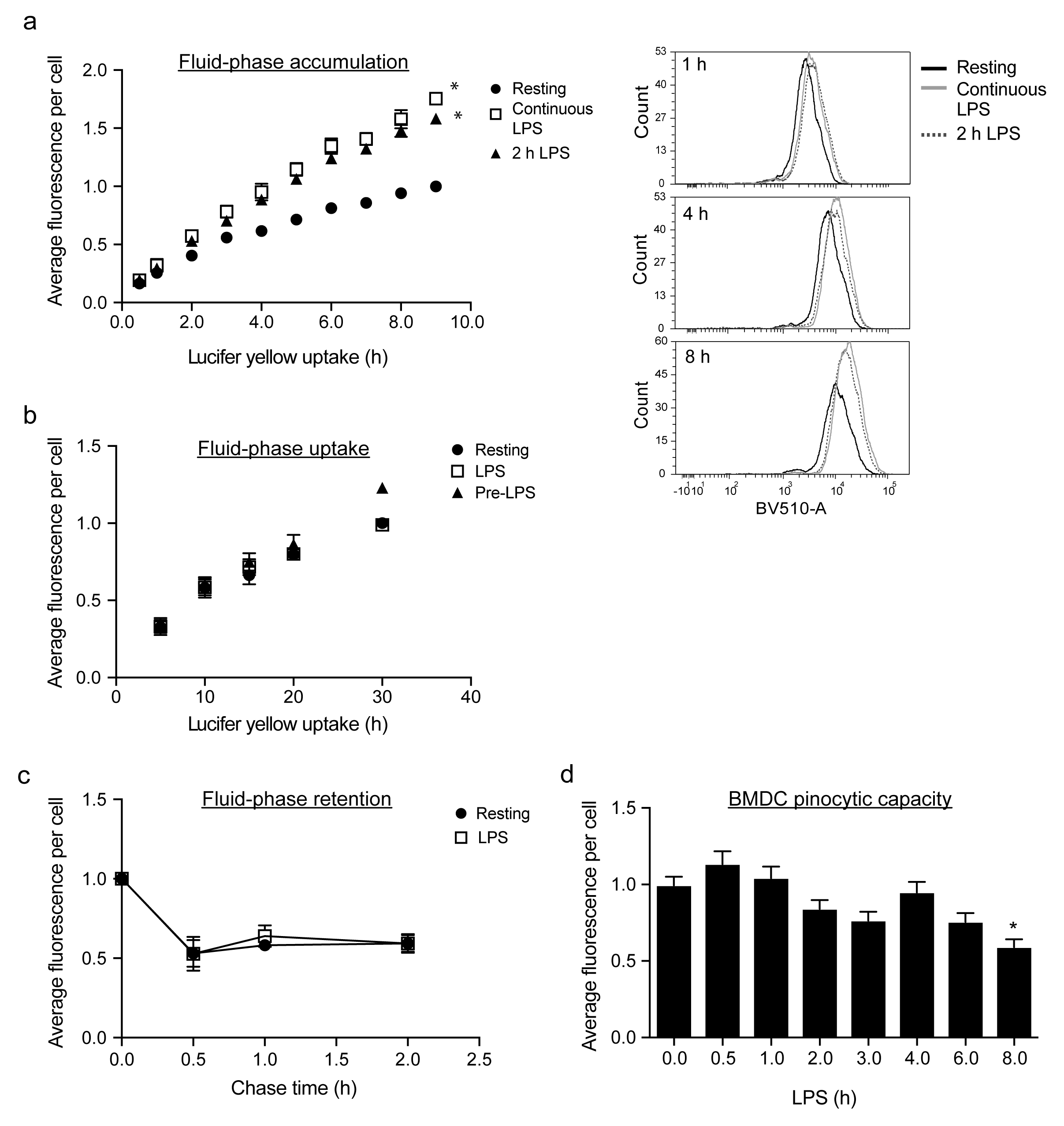

### S3 Fig

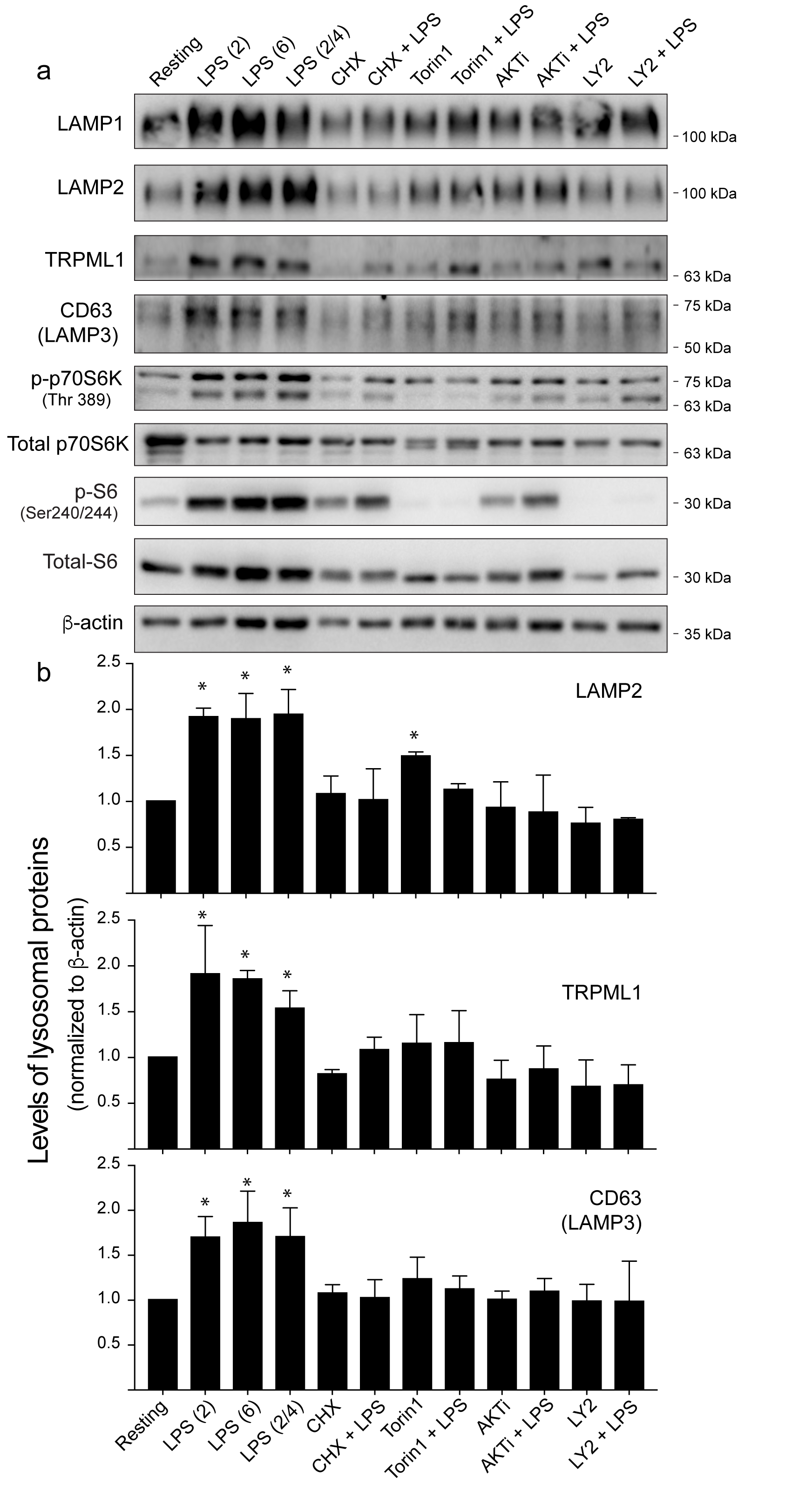

### S4 Fig

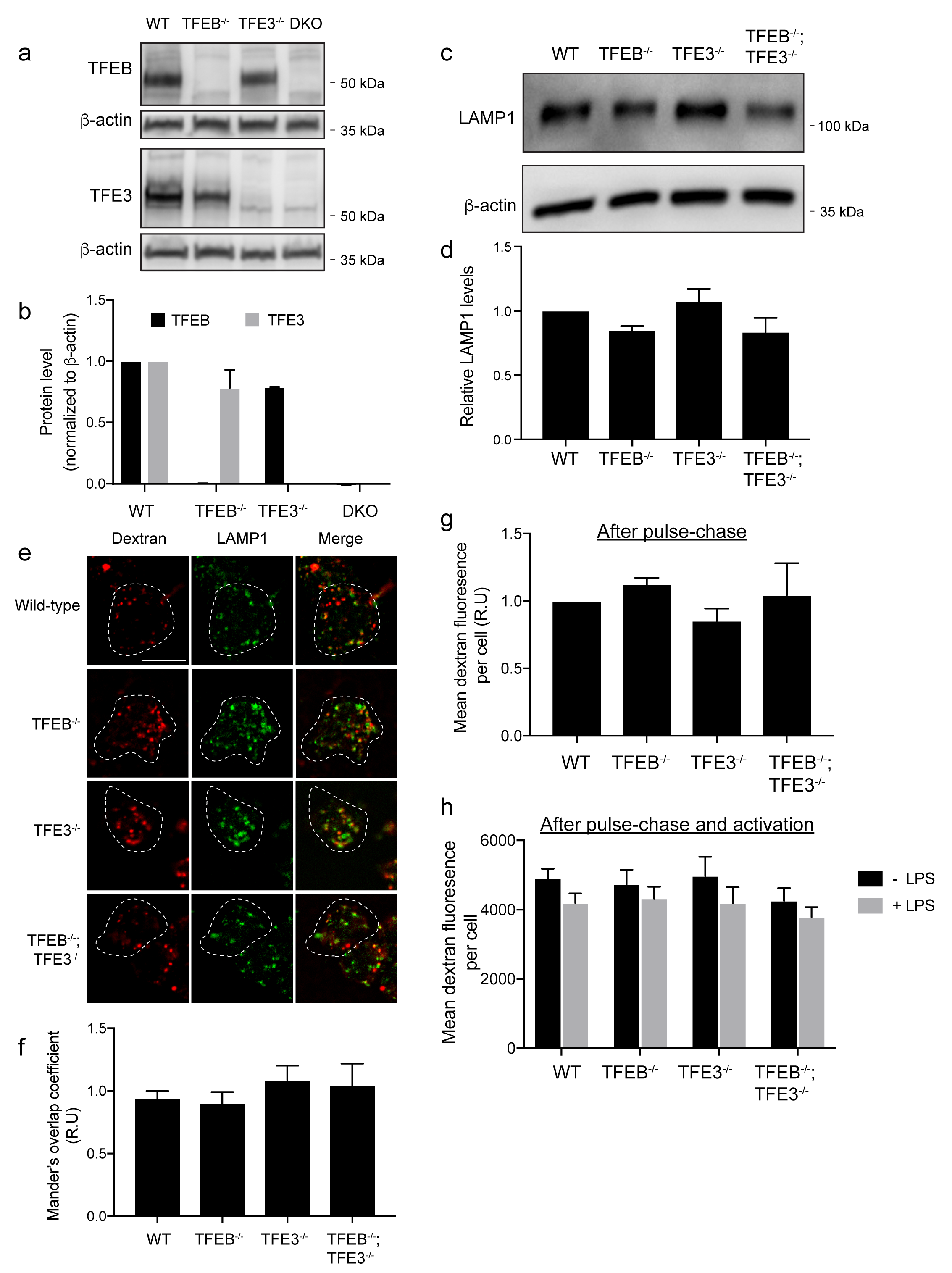

### S5 Fig

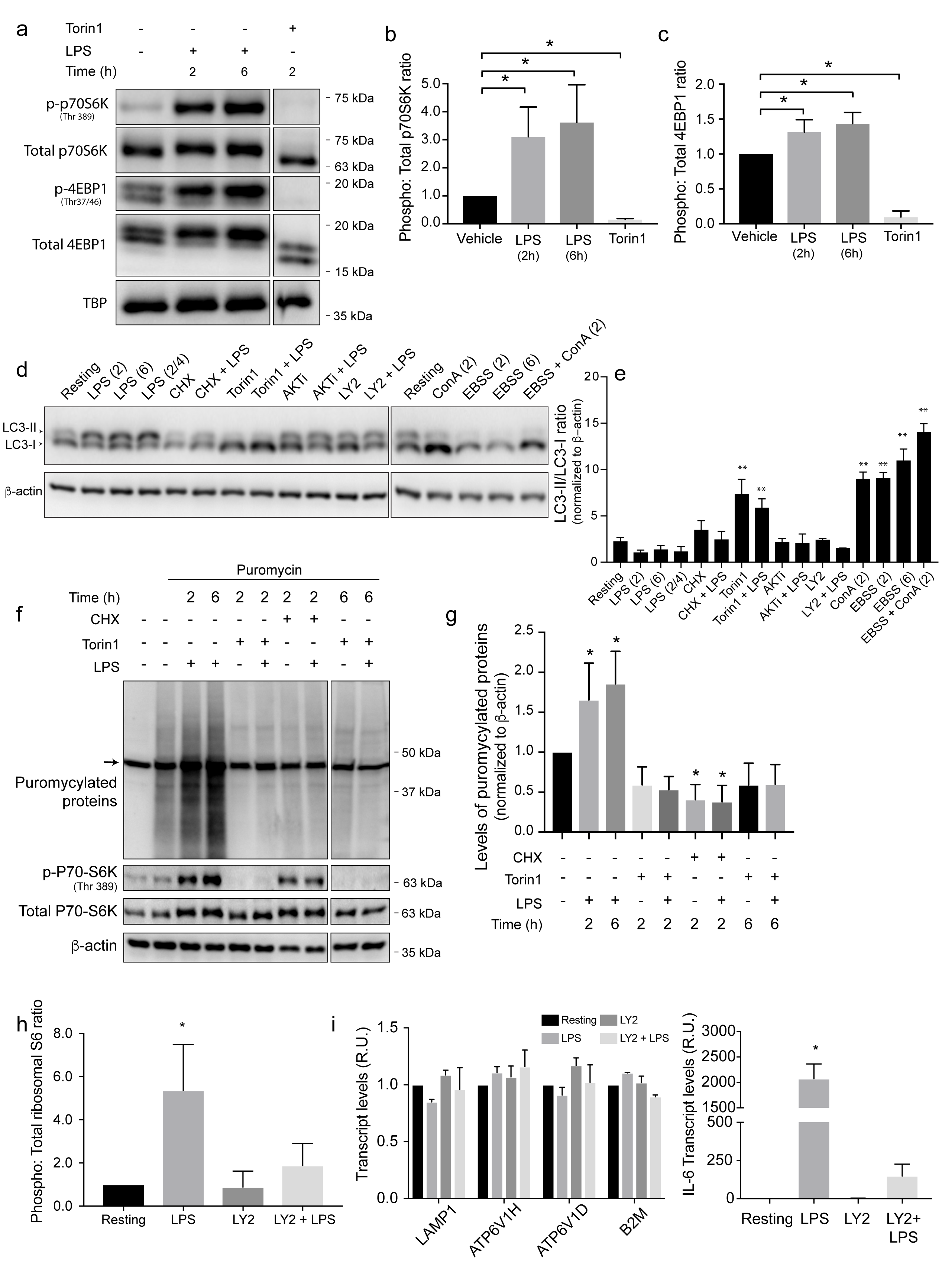

### S6 Fig.

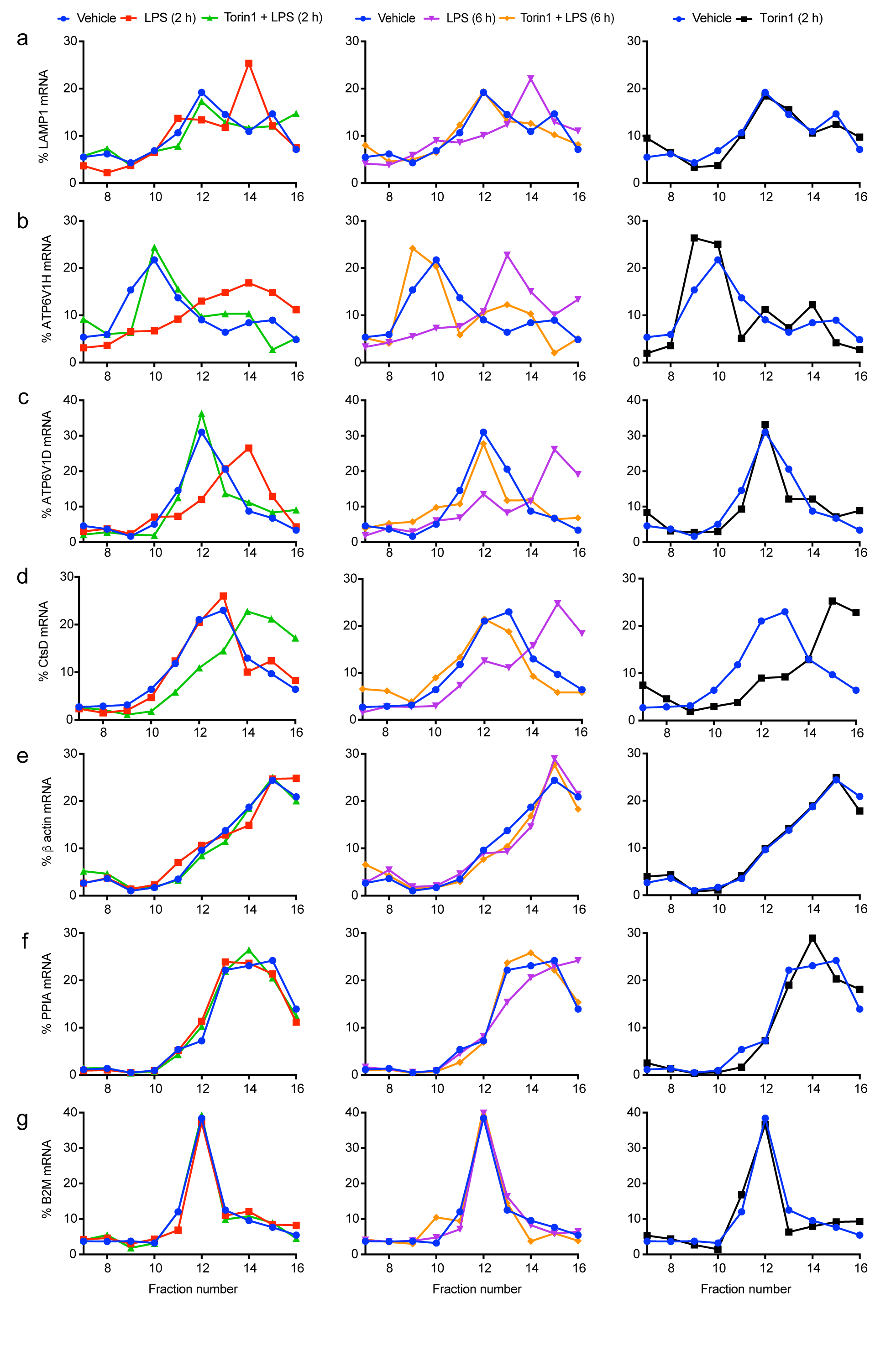

### S7 Fig.

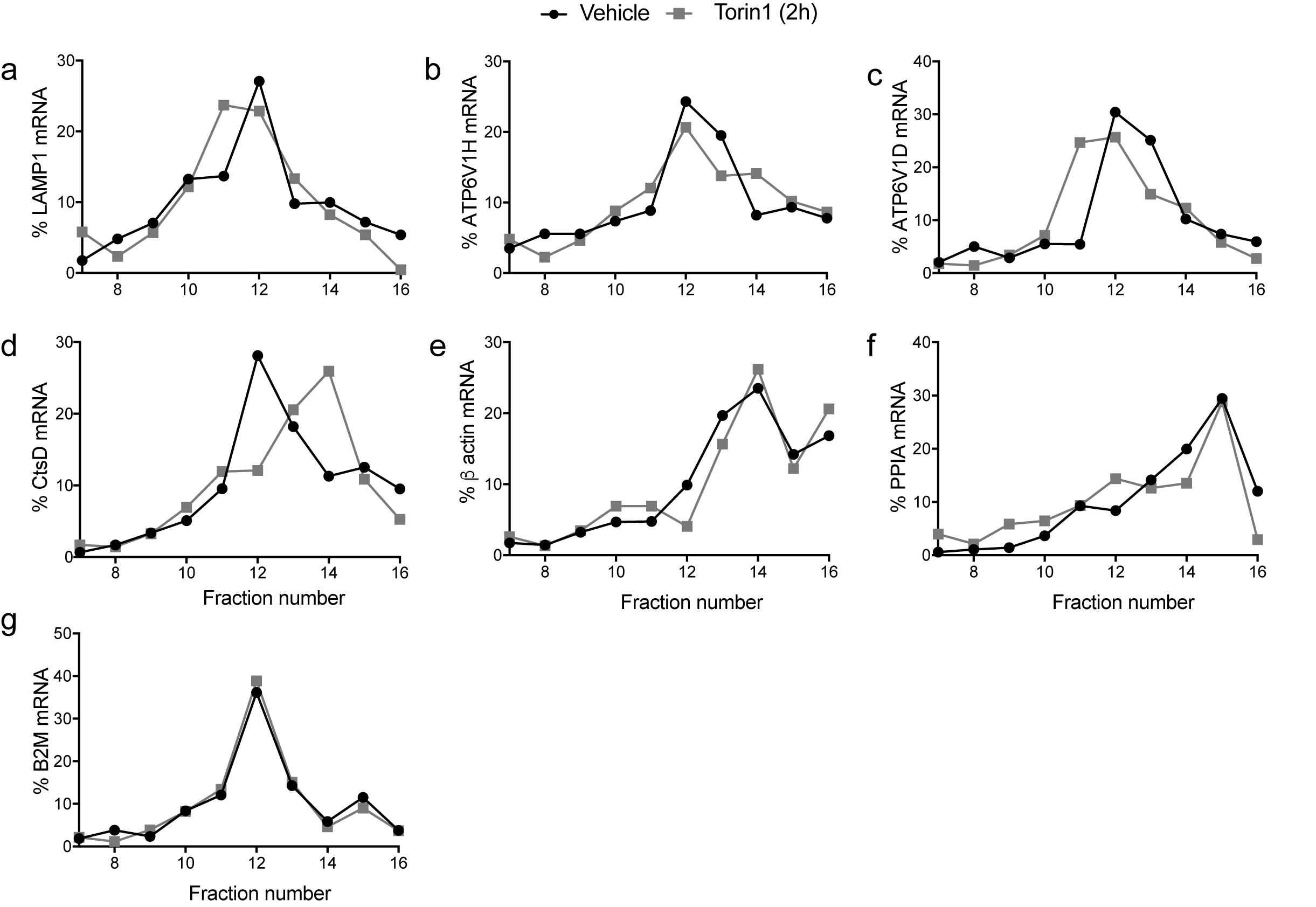

### S8 Fig.

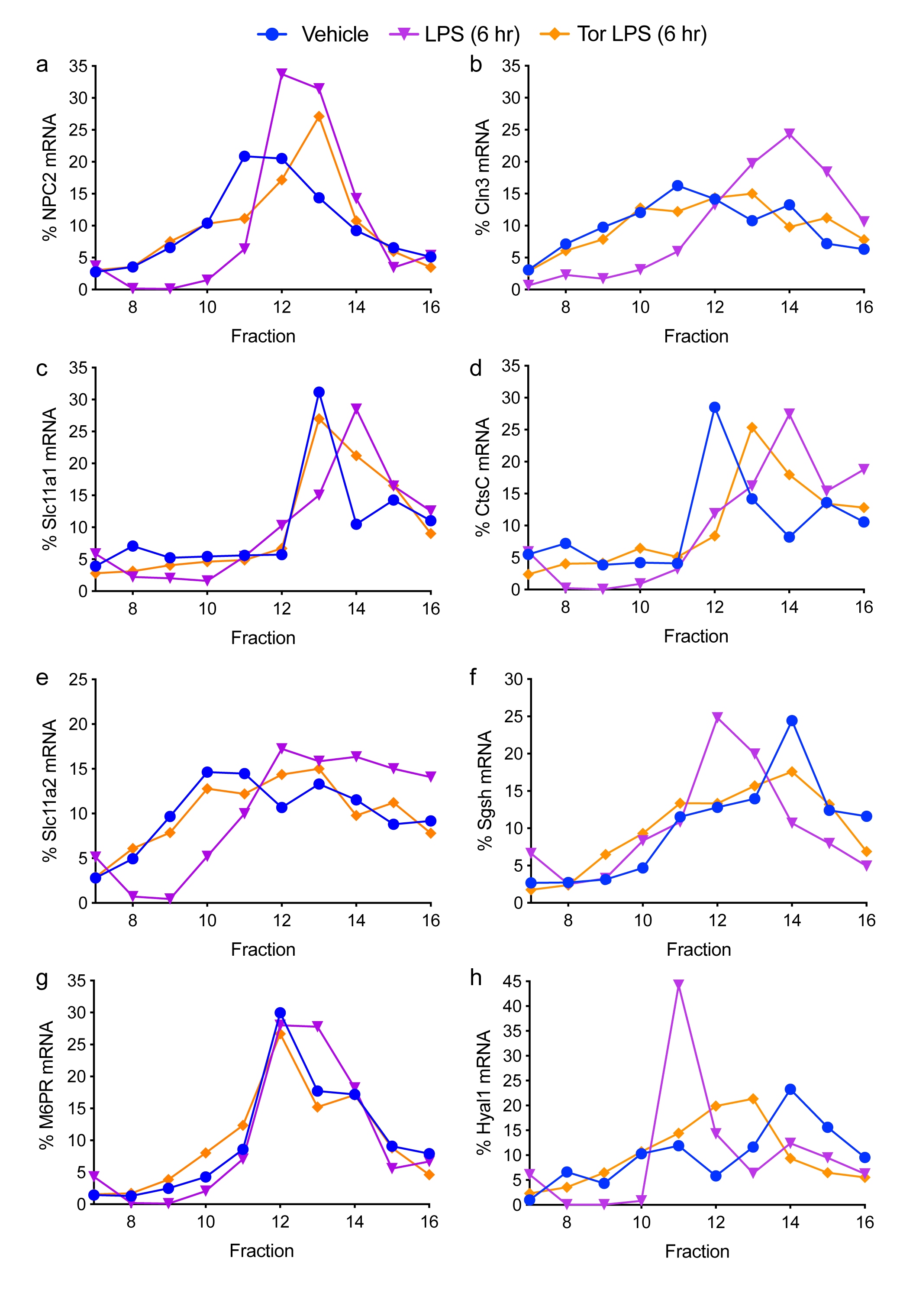

### S9 Fig.

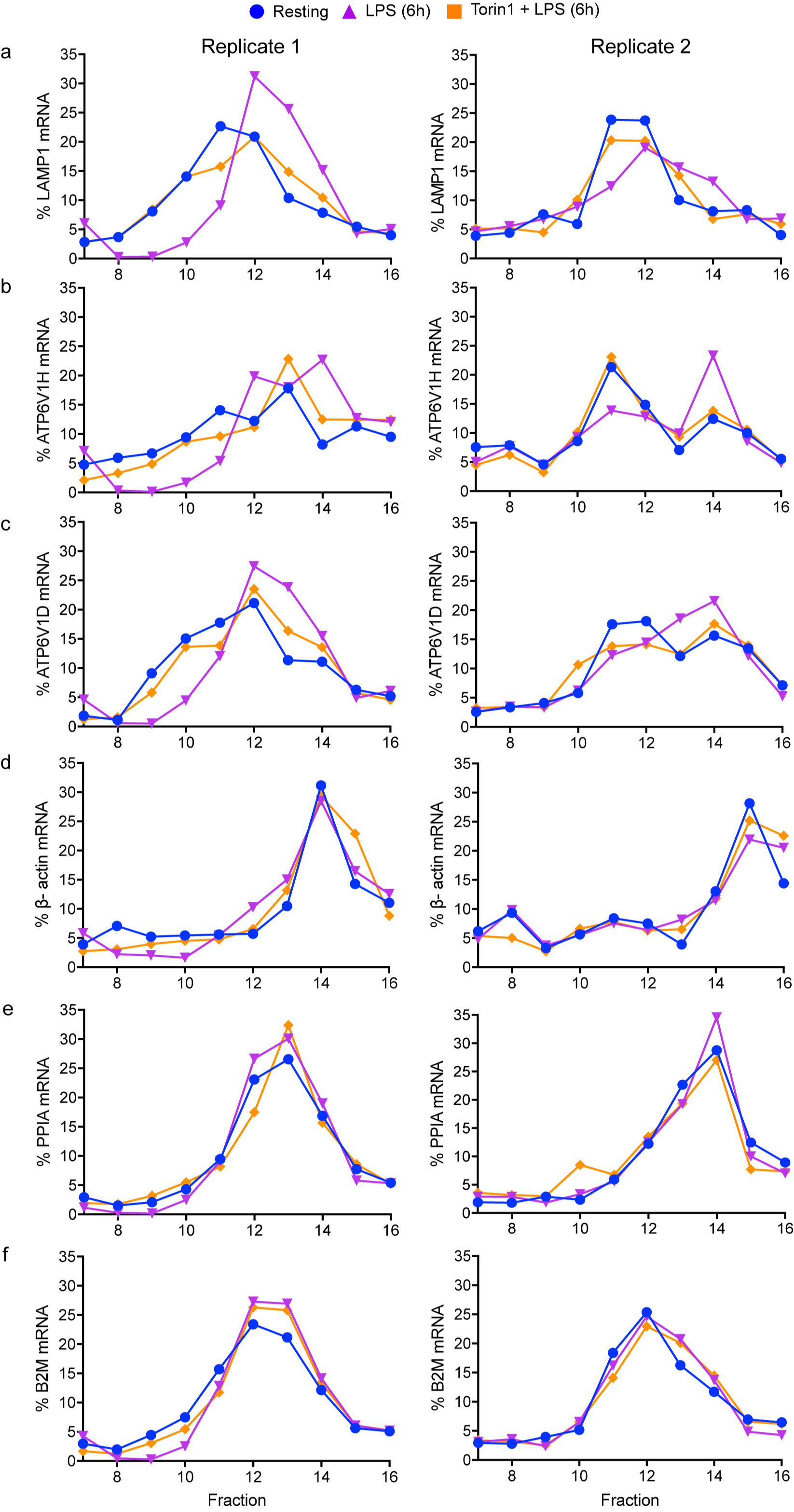

### S10 Fig.

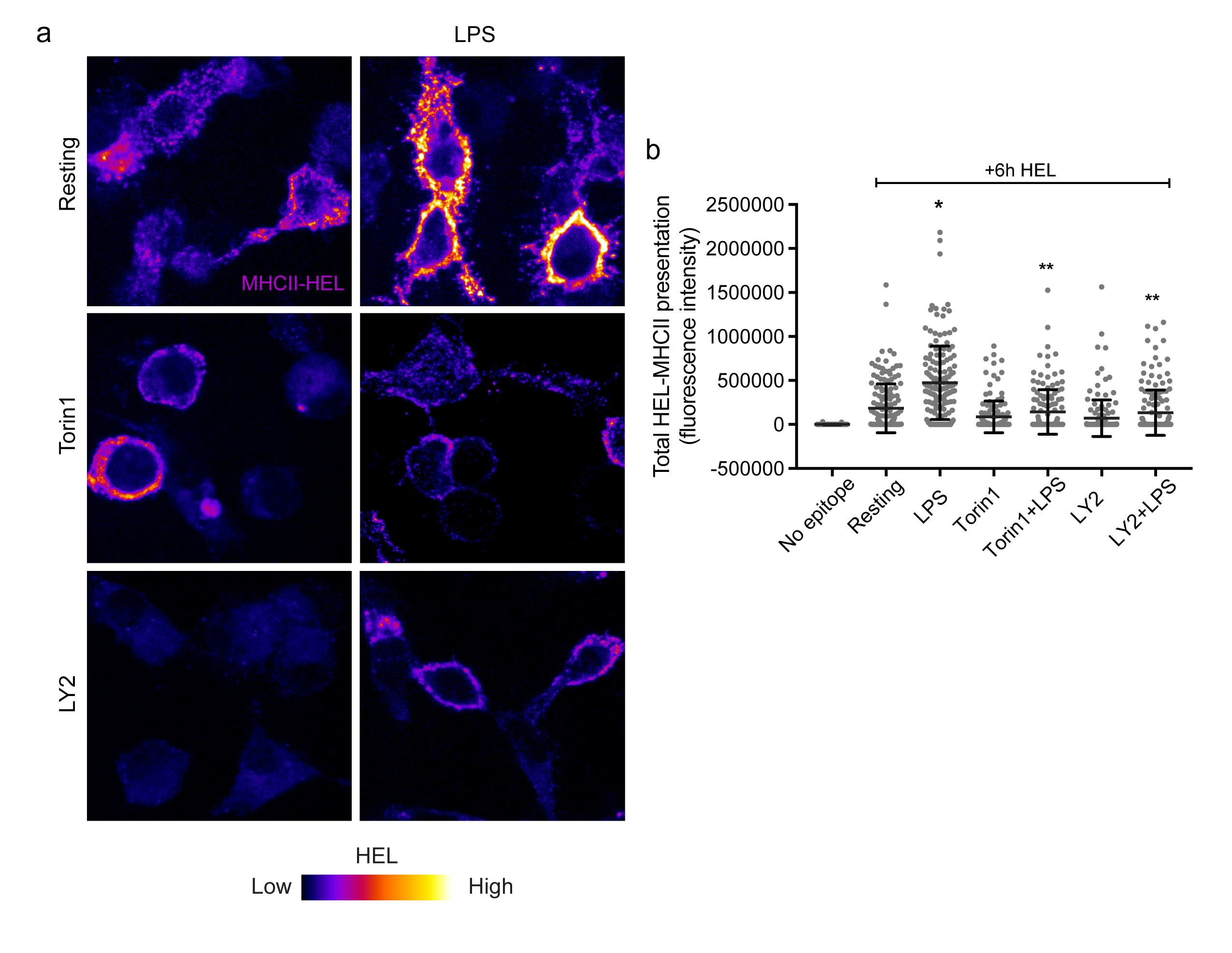

### S11 Fig.

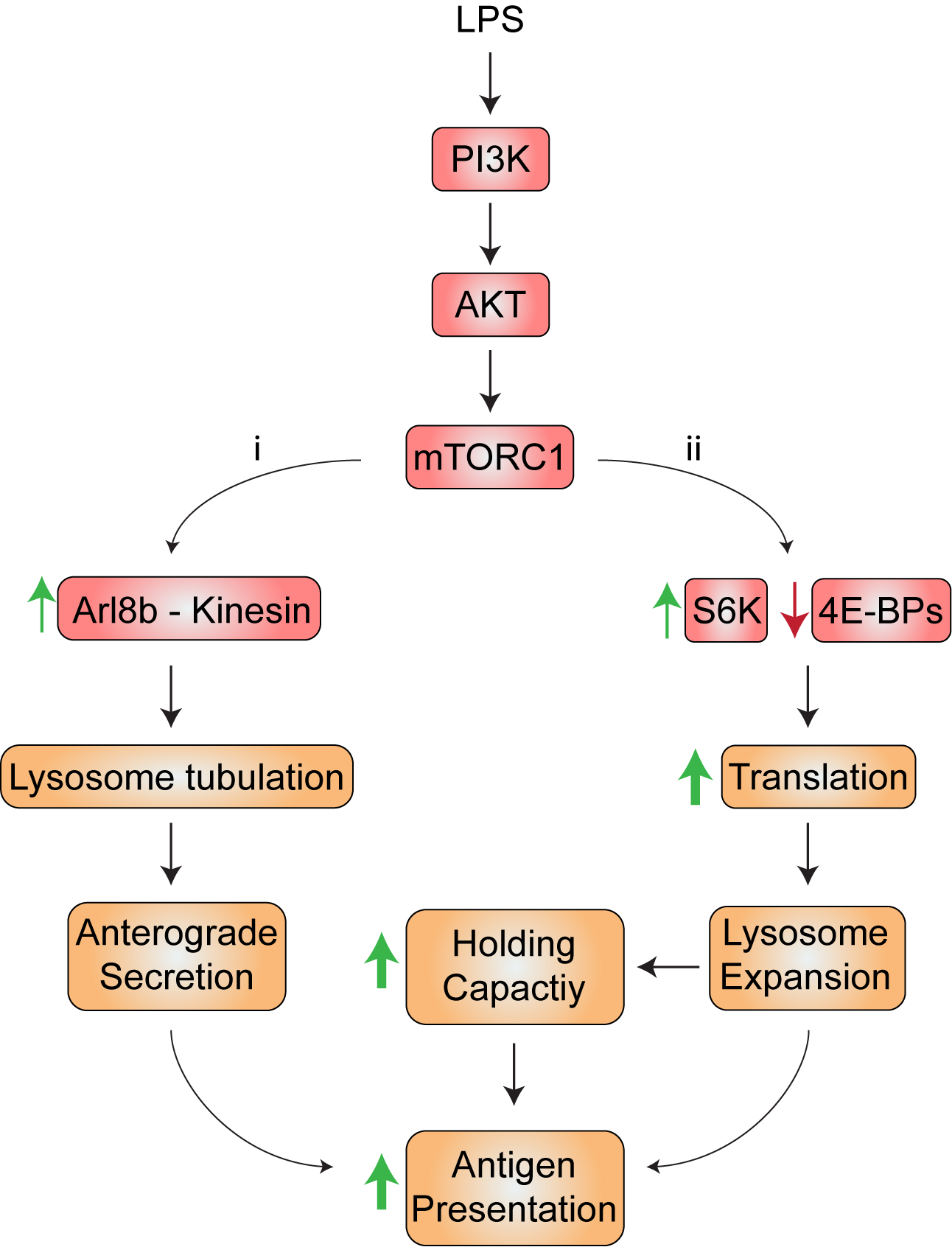
